# SP110 sequestration of SP100 protects against toxic filaments during innate immune signaling

**DOI:** 10.1101/2024.04.03.587867

**Authors:** Eric J. Aird, Julius Rabl, Tabea Knuesel, Lynn Scherpe, Daniel Boehringer, Jacob E. Corn

## Abstract

Stimulation of the innate immune system by foreign RNA elicits a potent response against invading pathogens and can trigger cell death. The mechanisms by which cells balance a robust response with cell-intrinsic lethality are still being uncovered. Employing genome-wide CRISPR-Cas9 genetic screens with triphosphorylated RNA stimulation, we identify speckled protein 110 (SP110) as a potent negative regulator of type 1 interferon-driven cell death. Death suppression by SP110 counteracts a death-promoting activity of another speckled protein, SP100. Both SP110 suppression and SP100 toxicity are mediated by direct interactions between the caspase activation and recruitment domains (CARDs) in each protein. SP100-induced death is mediated by homomeric CARD filaments that are disassembled by a heteromeric CARD interaction with SP110. Overexpression of SP100 is sufficient to overwhelm normal levels of SP110, leading to genotoxicity. Using cryo-EM and AlphaFold modeling, we develop and validate an atomic description of SP100 CARD filament formation and filament breaking by SP110. Genome-wide binding studies reveal that SP110 and SP100 normally associate at active promoters, but disruption of the CARD interaction releases SP100 to form toxic filaments. Overall, we uncover a novel regulatory partnership in human innate immunity that balances signal potency with cell intrinsic lethality.

## Introduction

The innate immune system guards against foreign pathogens and regulates cancer growth^1,2^. Both foreign and host molecules, including nucleic acids^3^, can stimulate an innate response and elicit the interferon signaling cascade. Interferon-stimulated genes (ISGs) perform downstream functions, including promoting the production of inflammatory cytokines and chemokines, resulting in further activation of both cell intrinsic and extrinsic responses^4^. Perturbing the innate immune response can trigger susceptibility to bacterial and viral infection, chronic inflammatory disease, or autoimmune disorders^5–7^. For example, knockout of ISG15, an interferon-stimulated negative regulator of type 1 interferon signaling, makes mice more susceptible to viral infection^8^. Conversely, loss of the interferon alpha/beta receptors (IFNAR1/2) stimulates growth and development of tumors^9,10^. An estimated 10 % of the human genome consists of ISGs, many of which are poorly characterized^4,11^.

Cell death is one of the intrinsic responses to innate immune signaling that cells use to combat infections to limit the replicative niche of invading pathogens^12^. Hyperactivation or alterations to innate immune signaling can disrupt the balance between sufficient innate immunity signaling and cell death. For example, the absence of key stress granule proteins G3BP1/2 results in excessive inflammation and cell death in response to cytosolic dsDNA^13^. Innate immune cell death frequently occurs through inflammasome assembly that leads to activation of caspases or physical disruption via pore-forming complexes^14^. However, the mechanisms responsible for maintaining homeostasis between an appropriate immune response and excessive cell death are incompletely described.

5’ triphosphorylated RNA (ppp-RNA) is a potent activator of the interferon response and ISG expression^15,16^. ppp-RNA, commonly found in bacteria and some viruses, is canonically recognized by the RIG-I signalosome^17^. However, many other proteins are reported to be involved in its recognition and response^18,19^. Interferon signaling induced by ppp-RNA leads to moderate amounts of intrinsic cell death^20,21^. Functional genomics has investigated factors that shape cellular fitness in response to various pathogens^22,23^; however, confounding factors such as cell entry and replication preclude drawing direct associations to molecularly defined innate immune activators such as ppp-RNA. A better understanding of the mechanistic roles of ISGs that balance the cellular response to innate immune stimulation by ppp-RNA with cell death could lead to novel biochemical insights and innovative approaches to treating infectious diseases.

Using multiple genome-wide CRISPR inhibition (CRISPRi) screens with ppp-RNA as a stimulus, we discover that speckled protein 110 (SP110) is a potent negative regulator of interferon signaling and downstream cell toxicity. This is mediated by an inhibitory direct interaction with SP100, whose upregulation is sufficient to trigger genotoxicity and cell death. A cryo-EM structure of SP100 reveals that it forms homomeric filaments via its caspase activation and recruitment domain (CARD), and these filaments are disassembled by interactions with the CARD of SP110. Point mutations disrupting the SP100 CARD filament prevent cell death, whereas mutations that impede SP110 heterodimerization promote interferon-induced toxicity. Our findings reveal a novel regulatory partnership between SP110 and SP100 that balances the strength of interferon signaling with cell death.

## Results

### SP110 loss confers hypersensitivity to interferon stimulation

Our lab and others recently reported that electroporation of triphosphorylated guide RNAs used in CRISPR genome editing elicits a remarkably strong interferon response^15,16^. Using a triphosphorylated guide RNA as a molecularly defined model for potent ppp-RNA stimulation, we sought to identify factors that regulate the balance between innate immune signaling and cell death. We employed a genome-wide CRISPRi screen to individually knock down (KD) expression of each human gene and assess the impact on cellular fitness upon ppp-RNA treatment (**Figure 1a**). The screen was performed in hTERT-RPE1 *TP53*^*-/-*^ cells, a non-cancerous and karyotypically normal cell line that has a robust interferon activation in response to foreign RNA^24^.

**Figure 1.**
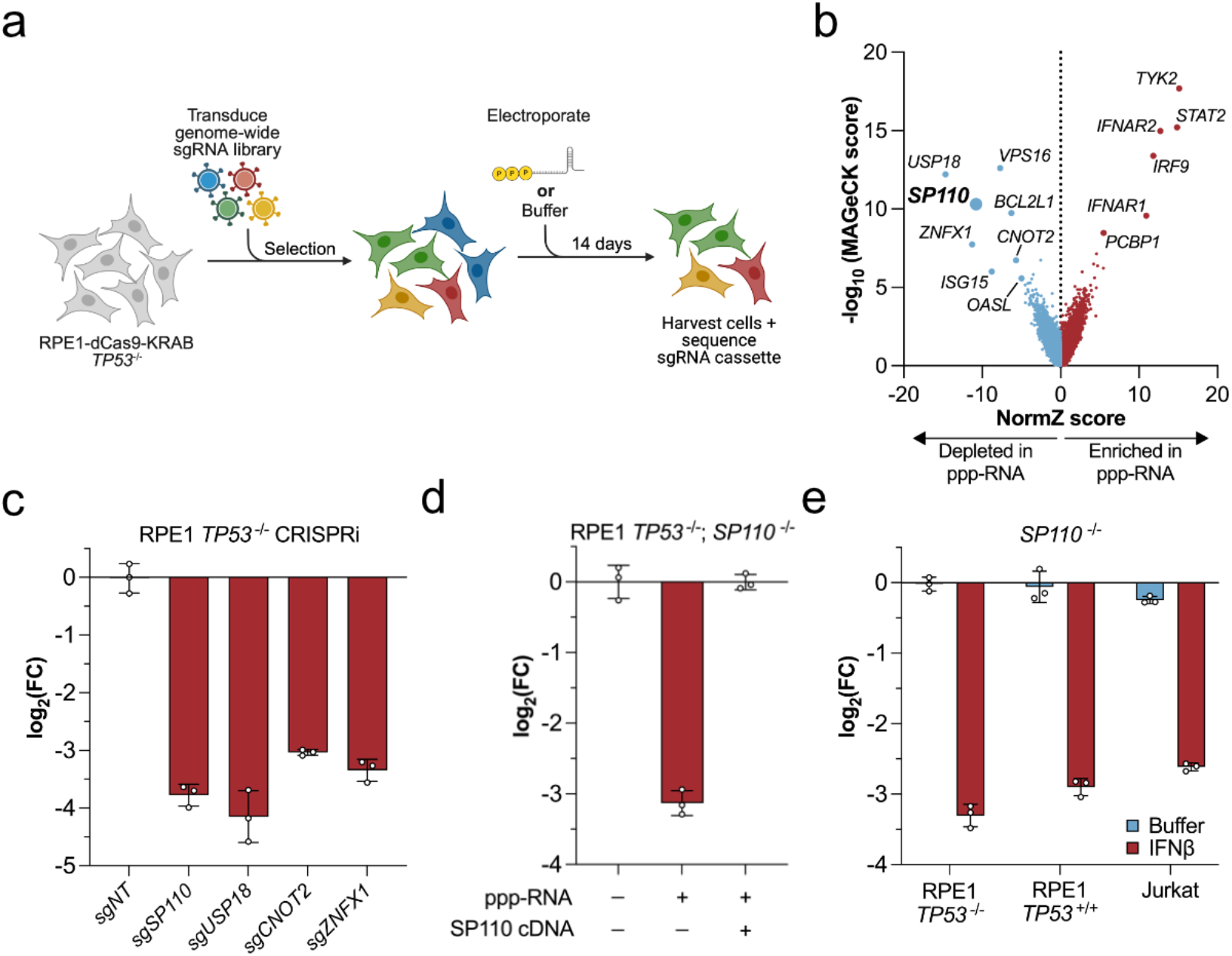
CRISPR screen reveals SP110 protects against hypersensitivity to interferon stimulation. **a**. Schematic for the genome-wide CRISPRi screen in RPE1 *WT* cells. **b**. Gene-level scores comparing the MAGeCK score versus NormZ score. Each data point represents a gene. Genes left of x = 0 were depleted in the ppp-RNA treated condition. **c**. Competition assay to individually validate candidate genes identified in the screen. All samples were treated with ppp-RNA. Data are plotted as the log_2_ value of the abundance of the indicated genotype relative to *WT* cells on day 7 versus day 0. sgNT = non-targeting sgRNA. **d**. Rescue experiment in *SP110* KO cells with full-length SP110 cDNA introduced via lentiviral overexpression. **e**. Competition assay in the indicated cell types in an *SP110* KO background either mock treated (Buffer) or with recombinant IFNβ. Each data point in **c, d**, and **e** represents an independent biological replicate. Bar graphs are displayed as mean +/- SD.

A combination of statistical metrics (see Methods) identified hundreds of genes that were significantly enriched or depleted in the ppp-RNA treated condition relative to mock treated cells (**Figure 1b, Supplementary Data 1**). Among these were several genes known to promote the transmission of the type 1 interferon response such as *IFNAR1/2, STAT2*, and *TYK2*. Guide RNAs targeting these genes were enriched in the ppp-RNA treated cells, presumably due to an inability to propagate an interferon response leading to a selective fitness advantage. RNA sensors of the DExD-box helicase (DDX) family such as *RIG-I* (*DDX58*) and *DDX56*^25^ were also enriched in treated cells, with the lower effect sizes reflecting their known redundancy in recognizing ppp-RNA. Conversely, negative regulators of the interferon response such as *USP18* and *ISG15* were significantly depleted in the treated condition. Gene set enrichment analysis (GSEA) reinforced that the screen accurately identified factors involved in innate immune signaling (**Figure S1a**).

Among the top depleted genes was SP110, one of four human SPs that are broadly linked to modulating the response to pathogen infections and regulating chromatin structure^26^. The role of SP110 in immunity is unclear since its presence promotes the persistence of certain viruses but knockout increases susceptibility to *M. tuberculosis* infections^27–29^. We sought to understand the function of SP110 in a pathogen-agnostic manner using ppp-RNA stimulated cell death as a model. We individually validated the top depleted genes from the screen using a growth competition assay and confirmed that *SP110* KD renders cells extraordinarily hypersensitive to ppp-RNA treatment, to a similar extent as known negative regulators such as *USP18* (**Figure 1c, S1b**,**c**). We also generated isogenic *SP110* knockout (KO) clones and verified that these phenocopy CRISPRi KD (**Figure S1d**). Hypersensitivity of *SP110* KO cells was rescued through complementation of full-length SP110 isoform C via lentiviral overexpression (**Figure 1d**), confirming the observed lethality is specific to loss of SP110.

We next tested whether SP110 was directly involved in RNA sensing but instead found that bypassing RNA recognition with recombinant interferon beta (IFNβ) addition also induced cell death in *SP110* KO cells (**Figure 1e**). We established that IFNβ hypersensitivity is neither *TP53* status nor cell type dependent as RPE1 *SP110*^-/-^; *TP53*^+/+^ cells and Jurkat *SP110*^-/-^ cells were hypersensitized to IFNβ (**Figure 1e, S1e**-**g**). Thus, SP110 is a potent negative regulator of interferon induced cell death that acts downstream of ppp-RNA sensing. The degree with which SP110 loss hypersensitizes cells to innate immune triggered cell death and the unclear mechanistic basis for this effect led us to further investigate how SP110 balances lethality with appropriate innate immunity signaling.

### SP110’s interactions with SP100 and PML bodies drives interferon susceptibility

To understand what protein(s) or pathway(s) promote interferon-stimulated toxicity upon SP110 loss, we performed a synthetic viability CRISPRi screen in RPE1 *SP110*^-/-^; *TP53*^*-/-*^ cells. As in wildtype (*WT*) RPE1 cells, *STAT2* and *TYK2* KD desensitized cells to ppp-RNA treatment, consistent with an SP110-independent effect (**Figure 2a, Supplementary Data 1**). However, we found that KD of *SP100* was strongly protective against ppp-RNA treatment in *SP110* KO cells without conferring a growth advantage to *WT* cells (**Figure 2a,b**). We further confirmed this insensitivity through generation of *SP110;SP100* dual KO cells (**Figure 2c, S2a**-**c**). SP110 and SP100 colocalize to phase separated nuclear promyelocytic leukemia (PML) bodies, and we furthermore observed that screen-based or individual KD of *PML* and other PML localized factors rescued the IFNβ hypersensitivity of *SP110* KO cells (**Figure 2a, S2d,e**).

**Figure 2.**
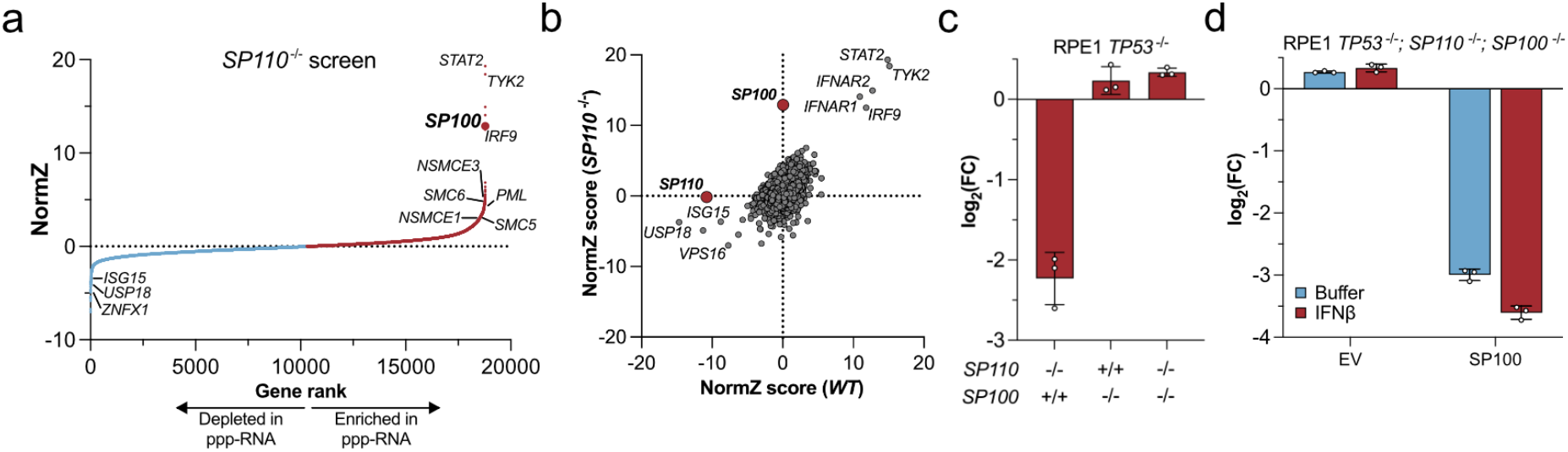
SP100 induces cellular toxicity in the absence of SP110. **a**. Ranked gene-level NormZ scores in the genome-wide screen performed in RPE1 *SP110* KO cells. Genes with y > 0 (red data points) were enriched in the ppp-RNA treated condition. **b**. Comparison of gene-level NormZ scores from genome-wide screens in *SP110* KO versus *WT* cells. **c**. Competition assay in the indicated genotype relative to *WT* cells treated with IFNβ. **d**. Rescue experiment in *SP110*;*SP100* dual KO cells with lentiviral overexpressed empty vector (EV) or SP100 treated with the indicated conditions. Each data point in **c** and **d** represents an independent biological replicate. Bar graphs are displayed as mean +/- SD.

We next performed SP100 rescue experiments in *SP110;SP100* dual KO cells. Reintroduction of SP100 isoform A in this background was sufficient to recapitulate interferon-induced cell death. Surprisingly, we also found that overexpression of SP100 was sufficient to induce cell death in the absence of IFNβ stimulation (**Figure 2d**). Even in *SP110* proficient cells, overexpression of SP100 induced lethality (**Figure S2f**). The magnitude of cell death in *WT* cells was lower than in *SP110;SP100* dual KO cells, implying the balance between SP100 and SP110 expression might be linked to the degree of toxicity. We explored this by titrating IFNβ in *SP110* KO cells and observed dose dependent toxicity (**Figure S2g**). We furthermore found that basal expression of *SP110* and *SP100* are highly correlated (r = 0.69) across 1,474 cell lines in the Cancer Cell Line Encyclopedia (CCLE)^30^ (**Figure S2h**). Indeed, *SP100*’s expression correlation with *SP110* across all lines in CCLE is higher than with any other transcript (**Figure S2i**). Taken together, our data suggest that SP110 prevents interferon-induced cell death by constraining a toxic activity of SP100.

### SP100 induces PML-dependent genotoxicity, senescence, and cell death

We hypothesized that SP100-induced cell death might be caused by a dysregulated interferon response. However, using RNA-seq to assess global transcriptomic changes in stimulated *WT, SP110* KO, and *SP110;SP100* dual KO cells, we observed very few significant changes in expression (56 genes with p_adj_ < 0.05) between the single and double knockout cells (**Figure S3a,b**). All three cell types mounted an equal interferon-induced transcriptional response as measured by upregulation of common ISGs (**Figure S3c**). The interferon-hypersensitive *SP110* KO cells showed no evidence of ISG hyperactivation, and the interferon-resistant *SP110*;*SP100* dual KO cells showed no evidence of ISG dampening. Transcriptome-wide, there was no interferon signature present among differentially expressed genes in either *SP110* KO or *SP110;SP100* dual KO cells when compared to *WT* cells (**Figure S3d,e**).

We next used immunofluorescence to visually inspect cell death in *SP110* KO cells after IFNβ stimulation. We observed marked micronucleus formation, a hallmark of genome instability, in over 60 % of *SP110* KO cells (**Figure 3a,b**). This was in comparison to < 2 % micronuclei in stimulated *WT* cells (**Figure S4a**). Since micronuclei frequently result from DNA damage, we stained for the marker 53BP1 and found that 67 % of *SP110* KO cells contained > 2 53BP1 foci compared to 15 % of *WT* cells in the presence of IFNβ (**Figure 3c**). ATM signaling, another sign of DNA damage, was upregulated in IFNβ treated *SP110* KO cells using phosphorylated KAP1 as a marker (**Figure 3d**). Notably, SP100 overexpression was sufficient to induce ATM signaling, even in the absence of IFNβ (**Figure S4b**). ATM signaling was not upregulated in the *SP110;SP100* dual KO cells, demonstrating the DNA damage is SP100-dependent. Loss of cyclin B1 and activation of p53 are linked with excessive DNA damage, cell cycle arrest, and the induction of cell death^31,32^. We detected an absence of cyclin B1 in interferon stimulated *SP110* KO cells (**Figure 3d**). Cyclin B1 loss was also observed when overexpressing SP100 (**Figure S4b**). We measured p53 activation using *CDKN1A* (p21) expression as a marker and again found upregulation only in *SP110* KO cells stimulated with IFNβ. No p21 upregulation was measured in *WT* or *SP110;SP100* dual KO cells, consistent with the observed DNA damage phenotype (**Figure 3e**).

**Figure 3.**
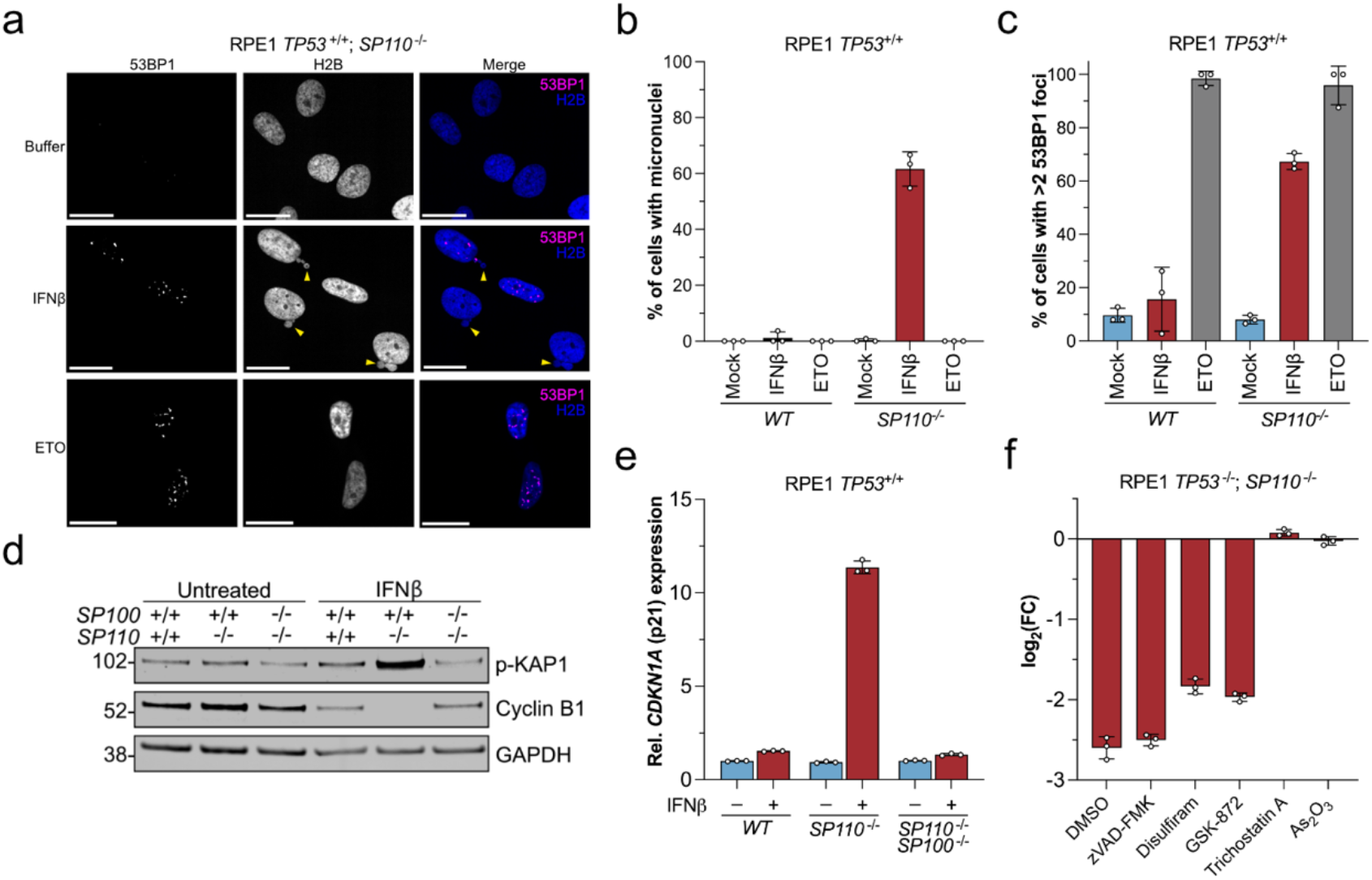
SP100 promotes genotoxicity, cellular senescence, and cell death. **a**. 53BP1 immunofluorescence in RPE1 *TP53*^+/+^ *SP110* KO cells stably expressing H2B-GFP treated with buffer, IFNβ (10 ng / mL), or etoposide (ETO; 25 μM) for 72 h. Scale bar = 20 μm. Yellow arrow denotes micronuclei. **b**. Quantification of micronuclei from **a** and **Figure S4a**. Each data point corresponds to > 50 cells from independent biological replicates. **c**. Quantification of cells containing > 2 53BP1 foci from **a** and **Figure S4a**. Each data point corresponds to > 50 cells from independent biological replicates. **d**. Western blot of whole cell lysates from untreated and 72 h IFNβ treated RPE1 *TP53*^+/+^ cells of the indicated genotype probing for phosphorylated KAP1 (S824) and cyclin B1 with GAPDH as the loading control. **e**. RT-qPCR of *CDKN1A* (p21) in untreated and IFNβ treated cells of the indicated genotype. Each data point corresponds to technical replicates from a representative experiment performed multiple times. **f**. Competition assay in IFNβ stimulated cells co-treated with the indicated compounds. Data are plotted as the log_2_ value of the abundance of *SP110* KO relative to *WT* cells at day 7 versus day 0. Each data point represents an individual biological replicate. Bars in **b, c, e**, and **f** are displayed as mean +/- SD.

Neither apoptosis, pyroptosis, nor necroptosis were individually responsible for SP100-induced lethality, as specific inhibition of each pathway did not substantially rescue cell death (**Figure 3f**). Therefore, a combination of programmed death pathways could result from excessive DNA damage. Completely blocking interferon signaling and perturbing SP110 and SP100 localization did prevent cell death. Trichostatin A, an HDAC inhibitor that prevents the expression of ISGs such as SP100 and SP110^33^, abrogated toxicity in IFNβ-stimulated *SP110* KO cells. And As_2_O_3_, a known disrupter of the SP100 / SP110-harboring PML bodies^34^, also prevented cell death. This latter finding corroborates our genetic perturbation data (**Figure S2**) and suggests that SP100 is specifically toxic when localized to PML bodies.

We observed significant colocalization of 53BP1 and PML foci in IFNβ treated *SP110* KO cells (r_median_ = 0.68), suggesting that SP100-induced DNA damage occurs in PML bodies (**Figure S5a**-**c,f**). Etoposide-treated cells had more 53BP1 foci than IFNβ treated *SP110* KO cells (**Figures 3c, S5d**-**f**), but less overlap between DNA damage and PML body localization (r_median_ = 0.53). In summary, we find that SP110 protects against interferon-induced SP100-dependent genome instability and cellular senescence in a PML body dependent manner, and that SP100 overexpression promotes this phenotype even in the absence of interferon.

### The SP110 CARD prevents SP100 CARD-induced lethality

To understand which molecular features of SP110 prevent interferon-induced cell death, we generated domain deletion cDNAs for stable lentiviral expression (**Figure 4a, S6a**). Like other SPs, SP110 contains a N-terminal CARD involved in protein-protein oligomerization and three C-terminal chromatin binding domains^26^. Though SP proteins are often associated with chromatin function^26^, we found that SP110’s chromatin binding domains were dispensable for cell protection since their deletion rescued hypersensitivity to IFNβ in *SP110* KO cells (**Figure 4b**). By contrast, an SP110 construct lacking only the CARD (βCARD) exhibited similar interferon hypersensitivity to the complete KO, and the CARD alone strikingly rescued cell death similar to full length. Confocal imaging revealed the CARD is required for the puncta-like localization of SP110, and we observed diffuse nuclear staining for βCARD (**Figure S6b**). The SP CARDs have high sequence homology^26^, but we found they do not functionally substitute for one another to prevent interferon-induced toxicity (**Figure 4c, S6c**). These data indicate a distinct role for the SP110 CARD in protecting from cell death.

**Figure 4.**
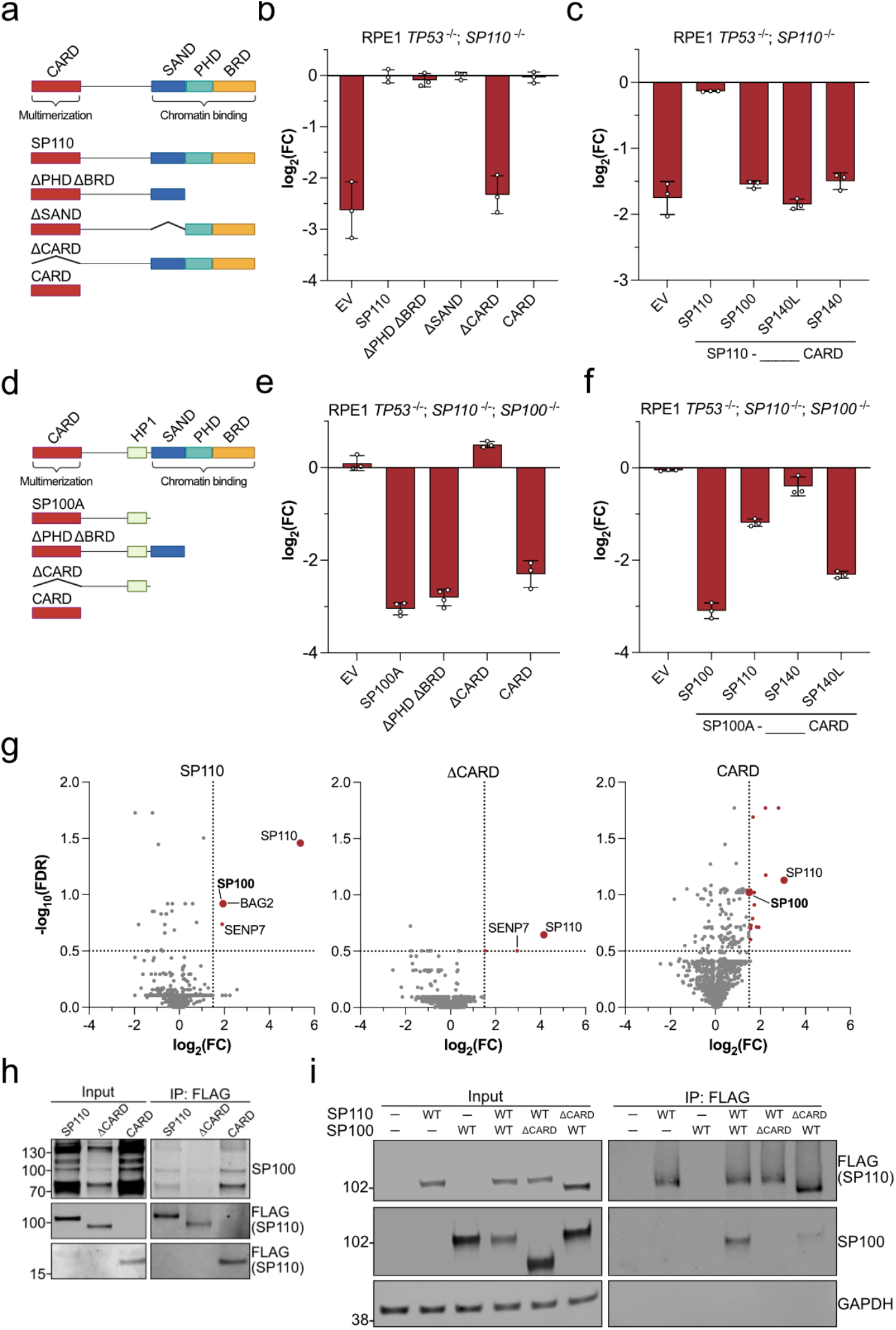
Physical CARD-CARD interaction between SP110 and SP100 prevents lethality. **a**. Schematic of SP110’s annotated domains. CARD = Caspase Activation and Recruitment Domain. SAND = Sp100, AIRE-1, NucP41/75, DEAF-1. PHD = Plant Homeobox Domain. BRD = Bromodomain. **b, c**. Rescue experiments of stably transduced SP110 domain mutants in *SP110* KO cells treated with IFNβ. In **c**, the SP110 CARD was replaced with the CARD from the indicated speckled protein. **d**. Schematic of SP100’s annotated domains. HP1 = Heterochromatin Protein 1. **e, f**. Rescue experiments of stably transduced mCherry-SP100 domain mutants in *SP110*;*SP100* dual KO cells treated with IFNβ. In **f**, the SP100 CARD was replaced with the CARD from the indicated speckled protein. **g**. Limma output of IP-MS spectral counts with the listed stably overexpressed FLAG-SP110 bait protein in *SP110* KO cells. Dotted lines indicate thresholds at x = 1.5 and y = 0.5. Proteins lying above both thresholds are labeled in red. **h**. FLAG co-IP in RPE1 cells of stably expressed FLAG-SP110 variants. **i**. FLAG co-IP in HEK-293T cells of transiently overexpressed full length (WT) or CARD deletion (ΔCARD) SP110 and SP100. Each data point in **b, c, e**, and **f** represents an independent biological replicate. Bar graphs are displayed as mean +/- SD.

We applied the same structure-function analysis to SP100 in *SP110;SP100* dual KO cells (**Figure 4d, S6d**). Overexpression of SP100 isoforms A (SP100A, βSAND βPHD βBRD) and B (βPHD βBRD), which lack chromatin binding domains, still caused marked cell death. Strikingly, the SP100 CARD alone was sufficient to cause cell death (**Figure 4e**), an exact inverse of the results with SP110 CARD. Toxicity was abrogated by removing the CARD from SP100 (βCARD). Like the SP110 CARD, SP100 localization to puncta-like foci requires the CARD (**Figure S6e**). We substituted other SP CARDs into SP100 and found they do not induce the same degree of toxicity as SP100’s CARD, except for SP140L (**Figure 4f, S6f**). The SP140L CARD has 85 % sequence homology due to a partial gene duplication of the SP100 CARD in primates^35^. In summary, SP100 toxicity only requires its CARD and is independent of its chromatin binding domains.

CARDs are often involved in protein-protein interactions to form homo- and hetero-oligomers^36^. We screened for interactors of SP110’s CARD using immunoprecipitation followed by mass spectrometry (IP-MS) with stably expressed full-length SP110, SP110 βCARD, and SP110 CARD alone in the *SP110* KO background. Only three proteins were significantly enriched with full-length SP110: SENP7, BAG2, and SP100 (**Figure 4g**). SENP7, a deSUMOlyase, was also enriched in ΔCARD but not in CARD alone, consistent with reported interactions with C-terminal SUMO motifs of SP110^37^. Strikingly, SP100 was enriched with full-length SP110 and CARD only, but not ΔCARD, suggesting a heteromeric CARD-CARD interaction. Notably, no other CARD-containing proteins, including other SPs or caspases, were enriched in any condition. We used co-IP and western blotting to validate both that SP110 and SP100 interact with one another, and that this interaction depends on the CARD from each protein (**Figure 4h,i**). Overall, our data reveal that the respective protective and cell killing activities of SP110 and SP100 are mediated by their CARD domains, that these proteins bind one another through a CARD-CARD interaction, and that loss of this interaction renders cells hypersensitive to interferon signaling.

### The SP100 CARD forms toxic filaments in the absence of SP110

SP110 and SP100 are both frequently associated with chromatin function^26^, so we assessed their localization on the genome. ChIP-seq for endogenous SP100 revealed that with a *WT* genotype, SP100 is predominately localized to promoters^38^ (**Figure 5a, S7a,b**). Surprisingly, we found in *SP110* KO cells that SP100 is no longer enriched at promoters, demonstrating that its localization depends upon SP110 (**Figure 5a, S7c**). We also observed minimal transcriptional changes between *SP110* KO and *SP110*;*SP100* dual KO cells, suggesting that SP100 has only a minor transcriptional role on its own (**Figure S3a**). We therefore investigated non-transcriptional functions of SP100 involving its CARD that might be triggered when it is freed from gene promoters in *SP110* KO cells.

**Figure 5.**
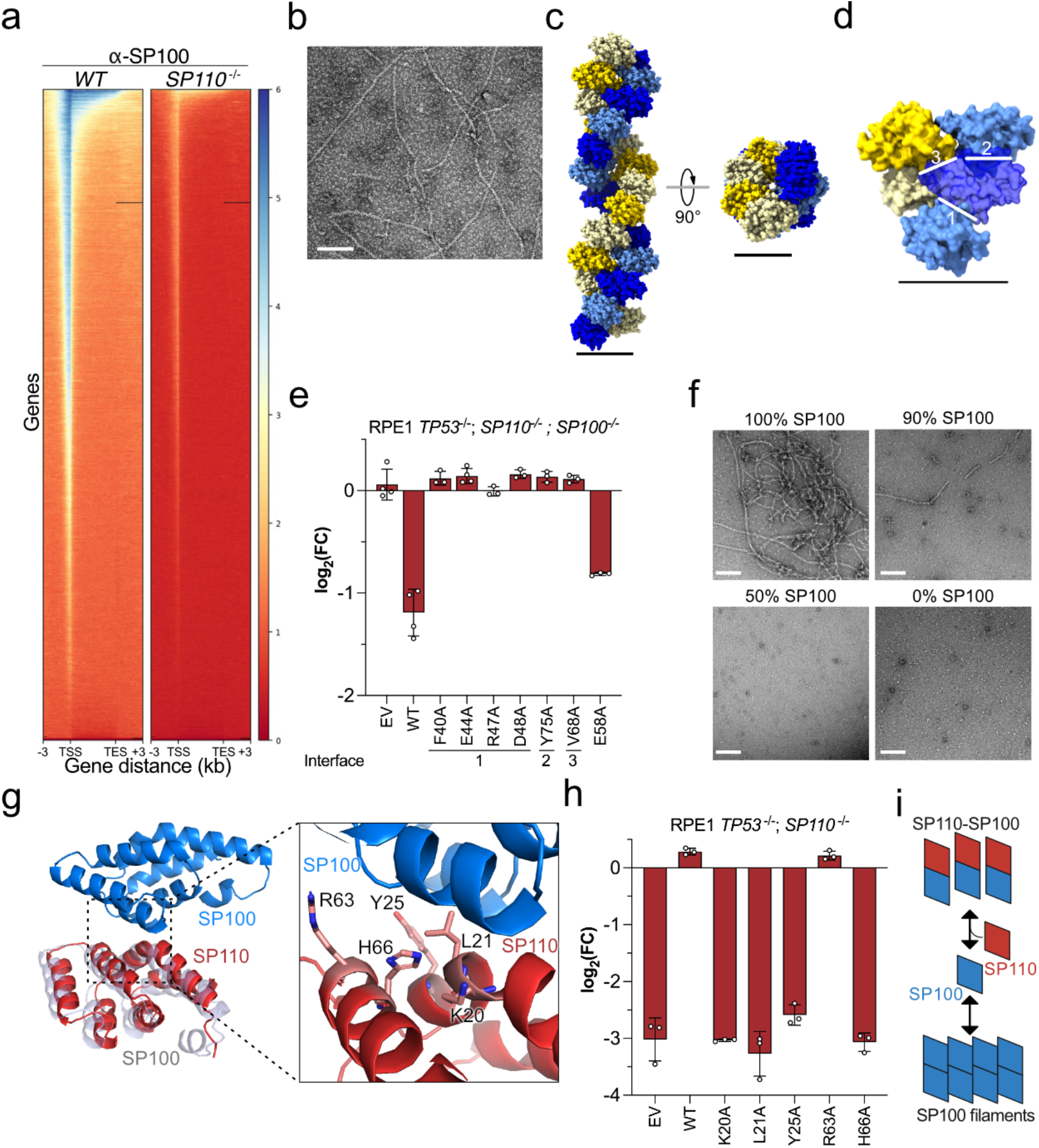
SP100 CARD filaments drive cellular toxicity. **a**. Heat map of endogenous SP100 ChIP-Seq peaks in and surrounding gene bodies of normalized length. Each line corresponds to a gene. The color scale indicates the relative intensity of the signal. **b**. Transmission electron microscopy (TEM) image of negative stained SP100 CARD filaments. Scale bar = 100 nm. **c**. Cryo-EM structure of the SP100 filament. Scale bar = 50 Å. **d**. Representation of CARD-CARD binding interfaces in the SP100 filament. The three major interfaces are indicated. Scale bar = 50 Å. **e**. Rescue experiments of stably transduced SP100-T2A-mCherry point mutants in *SP110*;*SP100* dual KO cells treated with IFNβ. WT = SP100. EV = Empty vector. **f**. TEM images of mixed SP100:SP110 at the indicated amount of SP100. Scale bar = 100 nm. **g**. AlphaFold2 model of a SP110-SP100 heterodimer. A displaced SP100 monomer is shown as a surface representation in grey. The inset highlights the SP110-SP100 CARD-CARD binding interface and the predicted important residues on SP110 are labelled. **h**. Rescue experiments of stably transduced SP110 point mutants in *SP110* KO cells treated with IFNβ. WT = SP110. **i**. Model for SP110 sequestration of SP100 to prevent filament formation. Each data point in **e** and **h** represents an independent biological replicate. Bar graphs are displayed as mean +/- SD.

CARD domains can form heteromeric interactions with other CARDs, but also homo-oligomeric filaments^39–41^. To explore the oligomeric state of the SP100 and SP110 CARDs, we recombinantly expressed and purified each CARD. Using negative staining and imaging by transmission electron microscopy (TEM), we observed that the SP100 CARD forms multi-micrometer long filaments (**Figure 5b**). These filaments provided the basis for acquiring and solving a 3.5 Å structure of the SP100 CARD in a homopolymer state using cryo-EM (**Figure S8, Table S1**). The SP100 CARD filament is comprised of two antiparallel strands that form a double helix (**Figure 5c**). The filament is 65 Å wide with a 10 Å central cavity and a twist length of 120 Å. This architecture stands in contrast to the three stranded CARD filaments of MAVS^39^ and NLRC4^40^, alluding to different functional niches (**Figure S9a**-**c**).

Each SP100 CARD, consisting of 6 alpha helices, has two interaction interfaces *in cis* on the same strand and one interaction interface *in trans* to the opposite strand (**Figure 5d**). We stably expressed SP100 with alanine point mutations at each binding interface in *SP110*;*SP100* dual KO cells. Each single mutation was sufficient to completely prevent SP100-induced lethality, whereas a mutation outside of the interface still induced cell death (**Figure 5e, S10a,b**). In summary, SP100 forms homo-oligomeric CARD filaments that induce cellular toxicity.

We observed no filament formation with recombinantly expressed SP110 CARD (**Figure S10c**). However, when testing the *in vitro* interaction between SP100 and SP110 CARDs, we found that SP110 had a striking ability to disassemble SP100 filaments. We mixed SP100 and SP110 at various ratios and imaged with TEM. As little as 1:10 SP110 CARD was sufficient to break SP100 filaments, and a 1:1 ratio led to a complete loss of filaments (**Figure 5f**). We used Alphafold2^42^ to model a SP110-SP100 CARD heterodimer, obtaining a strong interface predicted template modelling (ipTM) score of 0.86 (**Figure 5g**). This modelled heterodimer is predicted to block SP100 homo-oligomer formation and could be the molecular basis for SP110’s ability to inhibit SP100-induced cell death (**Figure 5g**). We tested this hypothesis by creating point mutations in full-length SP110. We found that four of five SP110 mutations at the putative interface with SP100 (interface #1) were sufficient to cause IFNβ-induced cell death, identical to complete *SP110* KO (**Figure 5h, S10d**). Taken together, these data reveal the molecular details of a functional CARD heteromeric interaction between SP110 and SP100 that inhibits formation of a toxic SP100 homomeric filament (**Figure 5i**).

## Discussion

Overall, we propose a model in which a balance between the interferon-stimulated factors SP110 and SP100 protects against interferon-induced lethality (**Figure 6a**). This is molecularly achieved by SP110 sequestering SP100 to gene promoters (**Figure 6b**). An imbalance in levels of SP110 leaves excess free SP100 which then forms genotoxic filaments. While we demonstrated this concept in a molecularly defined model system, the SP100 / SP110 regulatory partnership could have broader implications in the response to pathogens and in human disease.

**Figure 6.**
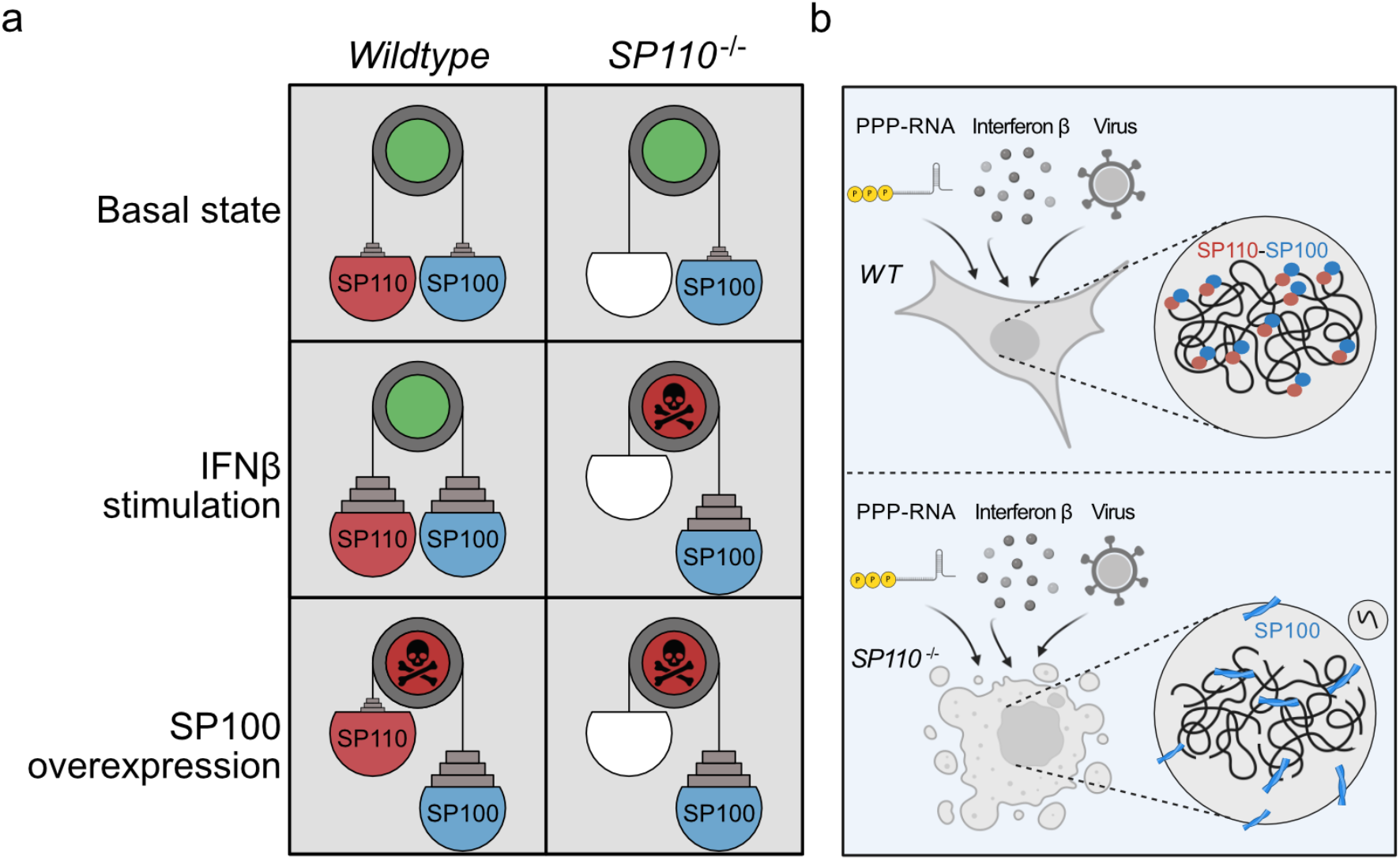
SP110 protects against SP100-induced genotoxicity. **a**. Schematic demonstrating the balance between SP110 and SP100 expression and the cellular outcome. **b**. Simplified model of the relationship between SP110 and SP100 in *WT* and *SP110* KO cells. Interferon stimulated *SP110* KO cells are unhealthy due to excess SP100 forming filaments, leading to DNA damage accumulation (broken chromatin) and micronuclei formation.

Human SPs share similar domain architecture yet perform distinct functions in maintaining immune homeostasis^26^. Here we establish a previously undescribed role of SP110 and SP100 in balancing an appropriate interferon response with cell death. In contrast, SP140 maintains heterochromatin status by repressing topoisomerases thus governing macrophage cell fate and response to bacterial challenges^43,44^. While SP140 expression is restricted to immune cells, SP110 and SP100 are expressed in nearly all cell types^45^. In mice, Sp140 negatively regulates interferon signaling, but Sp110 explicitly lacks this function^46^. However, mouse *Sp110* KO does not alter susceptibility to *M. tuberculosis* infection while mutations in human *SP110* increase susceptibility^46,47^. The three mouse and four human SPs share only ∼45 % sequence homology^26^, suggesting caution in extrapolating SP activities between organisms.

SP proteins, including S100 and SP110, are best known for their roles in chromatin transactions^26^. However, we find that the activity of SP110 and SP100 in interferon-stimulated cell death involves the CARD domain of both proteins in an unusual partnership. SP100’s cell killing capacity is achieved through formation of a CARD homo-oligomer that creates double-stranded helical filaments. Other innate immune factors form CARD-dependent oligomers, but these are distinct in both structure (**Figure S9**) and function^17,39,40^. AIRE, a close relative of SPs, forms homomultimers to facilitate transcriptional activity in T cells^48^. Recent reports indicate that AIRE is regulated *in cis* via its plant homeodomain (PHD) to promote polymerization on chromatin^49^. We instead find that SP100 polymerizes when not specifically localized on chromatin, and this is poisoned *in trans* by SP110 to block multimerization and sequester SP100 to gene promoters. It remains to be seen if other, unanticipated mechanisms exist to intentionally poison CARD oligomers found in other innate immune factors.

The functions of SP110 and SP100 parallel that of another ISG pair, PIM1 and GBP1. GBP1 forms needle-like structures that pierce Golgi bodies in the absence of PIM1-inhibitory phosphorylation, leading to necrotic cell death^50^. PIM1 thereby maintains GBP1 in a non-toxic state under basal conditions, much like SP110 restricts SP100-induced cell death. SP110 blocks SP100 filament formation through steric inhibition, whereas PIM1 chemically modifies GBP1 to prevent filament-like structures. While PIM1 / GBP1 activity occurs in the cytoplasm, SP110 / SP100 functions in the nucleus. By localizing to different cellular compartments, each protein pair occupies unique niches that could be applied to different innate immune contexts. Together with PIM1 / GBP1, stress granule proteins G3BP1/2^13^, and MORC3^51^, SP110 / SP100 represent a growing class of ISGs that regulate the balance between innate immune signalling and self-inflicted damage that limits pathogen spread and protects bystander cells.

## Methods

### Cloning, nucleotide, and protein sequences

All nucleic acid sequences can be found in **Table S2**. DNA was purchased from Integrated DNA Technologies (IDT) and Microsynth AG. *SP110* and *SP100* full-length and domain deletion constructs were amplified from cDNA generated from RNA isolated from RPE1 cells. Domain deletion constructs were assembled using site-directed mutagenesis. Other cloning procedures were performed using either Gibson assembly or golden gate assembly.

### Cell culture

hTERT-RPE1 (ATCC), hTERT-RPE1 *TP53*^-/-^ (generously provided by Dr. Steve Jackson), and HEK-293T (ATCC) cells were cultured in Dulbecco’s modified Eagle’s medium (DMEM, Gibco), supplemented with 10 % FBS and 100 μg / mL penicillin / streptomycin. Jurkat (ATCC) cells were cultured in RPMI medium (Gibco), 10 % FBS, and 100 μg / mL penicillin / streptomycin. All cells were incubated at 37 °C and 5 % CO_2_. Cell lines were routinely tested for mycoplasma contamination and tested negative (MycoAlert; Lonza). Cell lines were STR profiled (Microsynth AG). Recombinant human interferon beta (IFNβ; Abcam ab71475) was added at 10 ng / mL and incubated for 1-3 days. To induce DNA damage, cells were treated with etoposide (25 μM) for 24 h as a positive control. To modulate cell death pathways, zVAD-FMK (25 μM - apoptosis), disulfarim (10 μM – pyroptosis), or GSK872 (25 μM – necroptosis) were co-incubated with IFNβ treatment. Trichostatin A (500 nM) was used to inhibit ISG activation. As_2_O_3_ (10 μM) was used to disrupt PML bodies. Cell lines used in this study are listed in **Table S3**. All clonal cell lines were genotyped by Illumina sequencing and phenotyped for absence of protein production by western blot.

### Lentivirus production

Per reaction, 1 μg of genome vector, 800 ng of packaging plasmid (Addgene #8455; courtesy of Bob Weinberg), 200 ng of envelope plasmid (Addgene #8454; courtesy of Bob Weinberg), and 6 of μL polyethyleneimine (PEI) were mixed in 200 μl Opti-MEM and incubated for 10 min at RT. Each mixture was then added dropwise to a well in a 6 well-plate containing 5 x 10^5^ plated HEK-293T cells. After incubating for 2 to 3 days, the supernatant was collected, passed through a 0.2 μm filter, and used immediately or frozen at -80 °C. Cells were transduced in the presence of 8 μg / mL polybrene.

### In vitro transcription

ppp-RNA were produced in vitro using previously published methods^15^. Briefly, overlapping oligomers containing a T7 promoter, the desired protospacer, and sgRNA scaffold were amplified using Phusion polymerase (New England Biolabs). The unpurified DNA product was then subjected to in vitro transcription using the NEB HiScribe T7 High Yield RNA Synthesis Kit (New England Biolabs), incubating at 37 °C for 16 h. The following day, RNA was treated first with DNase I and purified with the miRNeasy mini kit (Qiagen), concentration measured by Nanodrop, and frozen at -80 °C.

### Electroporation

Electroporations were conducted in 96-well format (1 to 2 x 10^5^ cells per well) or with 100 μL cuvettes (5 x 10^6^ cells per cuvette) using Lonza 4D electroporation kits. The kit / program for each cell type was as follows: HEK-293T (SF kit / DG-130), Jurkat (SE kit / CL-120), and RPE1 (P3 kit / EA-104). After electroporation, 80 μL of prewarmed media was added to each cuvette and cells were allowed to sit at RT for 10 min before transferring to 2 mL medium in a 6 well-plate. For electroporation of Cas9-sgRNA to generate KOs, 100 pmol of Cas9 was complexed to 100 pmol of sgRNA, incubated for 10 min at RT, then added to cells / electroporation buffer mixture. For transfection of ppp-RNA, 50 pmol of *in vitro* transcribed RNA per 1 x 10^5^ cells was added to cells / electroporation buffer mixture.

### CRISPR inhibition screen

RPE1 *TP53*^*-/-*^ *WT* or *SP110*^-/-^ cells stably expressing dCas9-BFP-KRAB were transduced with a lentiviral CRISPRi sgRNA library^52^ containing ∼100,000 elements (5 sgRNAs per gene + non-targeting controls) at a target multiplicity of infection of 0.3 in the presence of 8 μg / mL polybrene. The lentivirus genome contained an untethered mCherry reporter gene to monitor transduction efficiency. 2 days post-transduction, cells were split into two replicates and treated with 10 μg / mL puromycin for 7 days. Cells were passaged and fresh puromycin added every 2 days. 9 days post-transduction, cells were electroporated with either *in vitro* transcribed ppp-RNA or a PBS buffer control. After roughly 10 doublings (13 days), cell pellets were collected and flash frozen. Genomic DNA was purified using the Puregene kit (Gentra), and the concentration was measured using a Nanodrop. The sgRNA cassette was amplified with NEBNext Ultra II Q5 HiFi polymerase (New England Biolabs) with indexed primers using the following steps with 5 μg of gDNA input per 50 μL reaction: 1x 98 °C for 30 s; 22x of 98 °C for 10s then 65 °C for 75 s; 1x 65 °C for 5 min. All common reactions were pooled and purified in a double SPRI bead purification (SeraMag; Cytiva). In the first round, 0.65X SPRI beads were added, and the supernatant collected. In the second round, 1X SPRI beads were used, and amplicon eluted in 20 μl H_2_O. The purified amplicons were analyzed on a TapeStation DNA flow cell (Agilent) for the expected amplicon size of 270 bp. Indexed amplicons were sequenced on a NextSeq 2000 P3 flow cell (Illumina) with 50 bp single end reads with a target of 50 M reads per sample. Reads were demultiplexed and analyzed by both MAGeCK^53^ and DrugZ^54^ first at the sgRNA level then the gene level to identify enriched and depleted genes. Gene-level ranks are listed in **Supplementary Data 1**. Gene set enrichment analysis (GSEA; v4.3.3) was performed separately on all enriched and depleted genes with reported gene lists having a nominative p-value < 0.05.

### Competition assay

Two populations of cells (*i*.*e. WT* versus *SP110*^-/-^ transduced with a fluorescent marker protein) were equally mixed on day -1 in a 6 well plate. On day 0, cells were analyzed by flow cytometry and treated with the indicated conditions. Cell populations were passaged and subsequently assessed by flow cytometry for 7 days post-treatment.

### Flow cytometry

Fluorescently labelled cells (via lentiviral transduction) were analyzed using an Attune NxT Flow Cytometer (Thermo Fisher), software v3.2.1. Data were analyzed in FlowJo (v10.9). The gating strategy is exemplified in **Figure S11**. Live cells were gated on SSC-A versus FSC-A followed by single cells on FSC-H versus FSC-A. Fluorophore presence was assessed by gating on SSC-A versus XFP (BFP, GFP, or mCherry).

### Clone genotyping

Genomic DNA was isolated using QuickExtract solution (Lucigen). Reactions were incubated for 10 min at 65 °C followed by 5 min at 98 °C and cooling to 4 °C. Primers containing Illumina adapter sequences were designed to amplify a ∼200 bp region surrounding the Cas9 target sites in *SP110* and *SP100*. Genomic DNA was amplified in two rounds using NEBNext Ultra II Q5 HiFi polymerase (New England Biolabs). In PCR 1, the above designed primers were used to amplify genomic DNA in 22 cycles. 1 μL from PCR 1 was used as input for PCR 2 where Illumina indexes were added in 10 cycles. Reactions were purified using 0.8x SPRI beads (SeraMag; Cytiva). Concentrations were measured on a Qubit and analyzed on a TapeStation DNA flow cell (Agilent). Samples were sequenced on either a MiSeq or NextSeq2000 with 2 x 150 paired end reads. The target read count per sample was 100,000 reads. Reads were demultiplexed and analyzed with CRISPResso2^55^ (v2.0.20b) using default settings.

### Western blot

Cell lysates were collected by pelleting, washing, and resuspending cells in 1x RIPA buffer supplemented with 1x protease inhibitor cocktail (Thermo Fisher). Lysates were incubated on ice for 15 min, pelleted at 15k x g for 15 min, and the soluble fraction collected. Protein concentration was determined using Bradford assay. 20 μg of input was electrophoresed on a 4-20% SDS-PAGE gel. A wet transfer to a 0.8 μm nitrocellulose membrane was performed at 4 °C. The membrane was blocked in TBS-T + 5 % milk for 45 min followed by overnight incubation at 4 °C in TBS-T + 2% BSA containing primary antibody. The antibodies and concentrations used are found in **Table S4**. The next day, membranes were washed in TBS-T and then incubated in TBS-T + 5 % milk + 1:10,000 secondary antibody (LI-COR 800CW and / or 680RD). Membranes were again washed in TBS-T. A final wash in PBS was performed prior to imaging on a LI-COR Odyssey using the 800 and 700 nm channels.

### Reverse transcription quantitative PCR (RT-qPCR)

Cellular RNA was isolated using the RNeasy kit (Qiagen) with on-column DNase digestion. RNA concentration was measured by Nanodrop, and 125 ng was used as input for reverse transcription (iScript cDNA Synthesis Kit; Bio-Rad). qPCR was then performed with a 1:10 dilution of cDNA using SsoAdvanced Universal SYBR Green Supermix (Bio-Rad) on a QuantStudio 6 (Thermo Fisher) per manufacturer’s protocol. Data were then analyzed by the ddCt method using *ACTB* as the endogenous control.

### RNA-seq

1 x 10^6^ RPE1 *WT, SP110*^-/-^, or *SP110*^-/-^; *SP100*^-/-^ cells were electroporated +/- ppp-RNA and collected 6 h later. Cellular RNA was isolated with the RNeasy kit (Qiagen) with on-column DNase digestion. RNA samples were sent to Novogene for library preparation and sequencing. Experiment was performed with three biological replicates. Reads were aligned to the genome using STAR^56^, and differentially expressed genes were annotated using DEseq2^57^. The DEseq2 results are given in **Supplementary Data 2**.

### Immunofluorescence

Stably expressing H2B-GFP RPE1 *WT* and *SP110*^-/-^ cells were generated using lentivirus generated from Addgene plasmid #91788 (Courtesy of Beverly Torok-Storb). Cells were plated on 8 well glass slides (Ibidi) and incubated in +/- 10 ng IFNβ for 3 days. Cells were fixed in 4 % PFA for 15 min at RT, washed in PBS, and permeabilized in PBS + 0.1 % Triton X-100 for 15 min. Following another PBS wash, cells were blocked in PBS + 2 % BSA for 1 h at RT. Primary antibody (**Table S4**) was added in PBS + 1 % BSA and incubated overnight at 4 °C. The next day, cells were washed and incubated in PBS + 1% BSA with fluorophore-conjugated secondary antibody for 1 h at RT. Cells were washed, air dried, and mounted in ProLong Diamond Mountant (Thermo Fisher). Slides were imaged on a Nikon Spinning Disk SoRa (ETHZ ScopeM) with a 100x oil-immersion objective. Images were analyzed using ImageJ^58^ (v2.14.0). For 53BP1 foci quantification, images had a constant threshold applied, and then the “analyze particles” function was used. Micronuclei were counted manually in the H2B channel. PML and 53BP1 Pearson object correlation was calculated using the JACoP plugin^59^.

### Immunoprecipitation mass spectrometry (IP-MS)

RPE1 *SP110*^-/-^ cells were stably transduced with FLAG-tagged SP110, SP110 ΔCARD, or SP110 CARD and selected with puromycin for 7 days. The input for immunoprecipitation was harvested from 5 x 10^6^ cells stimulated for 24 h with 10 ng / mL IFNβ. All lysis steps were performed on ice. Cells were pelleted, washed, and resuspended in cytoplasm lysis buffer (10 mM Tris-HCl, pH 8, 150 mM NaCl, 320 mM sucrose, 0.5 % Igepal, 0.1 mM EDTA, protease inhibitor cocktail) for 15 min. The nuclei were gently pelleted and resuspended in nuclear lysis buffer (50 mM Tris-HCl, pH 8, 500 mM NaCl, 1 % Igepal, 10 % glycerol, 1 U / mL benzonase, 1 mM MgCl_2_, protease inhibitor cocktail). The nuclei were lysed by passing through a douncer 30 times. The insoluble fraction was pelleted, and the supernatant used as input for immunoprecipitation with anti-FLAG M2 magnetic beads (Sigma). Beads were incubated overnight at 4 °C, rotating. Beads were washed in 1x TBS, and then bait protein was eluted by incubating in TBS containing 150 ng / μL FLAG peptide for 10 min, shaking. Samples were submitted to the mass spectrometry facility at the Functional Genomics Center Zurich. Data were analyzed using Limma^60^ using spectral counts as the input. The Limma output is given in **Supplementary Data 3**.

### Coimmunoprecipitation (Co-IP)

2 x 10^5^ HEK-293T cells per condition were transiently transfected with 1000 ng of each plasmid in the presence of PEI. 48 h later, cells were collected and lysed directly in nuclear lysis buffer (above). Lysates were clarified and used as input for immunoprecipitation with anti-FLAG M2 magnetic beads. The input and elution fractions were analyzed by western blot.

### ChIP-Seq

10 x 10^6^ RPE1 *WT* or *SP110*^-/-^ cells were incubated with 10 ng / mL IFNβ for 6 h. Cells were detached, washed, fixed in 1 % PFA for 15 min at RT, and quenched with 50 mM glycine. Cell pellets were washed, and flash frozen in liquid nitrogen. Pellets were lysed in 3 sequential lysis buffers to remove cytoplasm and obtain isolated nuclei according to a previously published protocol^61^. Nuclei were sonicated with a Covaris S220 to obtain a desired fragment length of 300 bp. Lysates were then incubated overnight at 4 °C with 5 μg of prebound anti-SP100 or rabbit IgG isotype control protein A dynabeads (Thermo Fisher). The next morning, beads were washed in 6x RIPA buffer followed by 1x TBS and eluted in 50 mM Tris-HCl, pH 8, 1 mM EDTA, 1 % SDS by incubating at 65 °C overnight to reserve crosslinks. The following day, the dynabeads were collected on a magnetic stand and the supernatant was collected. The supernatant was then incubated with RNase A and proteinase K (Qiagen). DNA was purified (MinElute kit, Qiagen) and analyzed on a TapeStation (Agilent). The NEBNextII kit (New England Biolabs) was used to prepare the sequencing library, and the indexed library was sequenced on a NextSeq 2000 P3 flow cell (Illumina) with 50 bp single end reads with a target of 25 M reads per sample. Reads were demultiplexed, aligned with Bowtie2^62^, and peaks called with MACS3^63^. Graphics were generated using deepTools^64^ and ChIPseeker^65^ (v1.32.1).

### Recombinant protein production

Human SP110 and SP100 CARD domain cDNA sequences were cloned into pET28b backbone with an N-terminal His_6_ tag, maltose-binding protein (MBP), and TEV cleavage site (**Table S5**). Each construct was transformed into BL21 Rosetta 2 (Merck) and grown in LB at 30 °C. At an OD_600_ of 0.6, the cells were transferred to 18 °C for 30 min, then IPTG was added to a final concentration of 0.5 mM to induce expression. Cells were incubated for 16 h at 18 °C then pelleted at 4,000 x g for 30 min. The cell pellet was lysed in lysis buffer (20 mM HEPES, pH 8, 200 mM NaCl, 10 % glycerol, 5 mM imidazole, 1 mM PMSF) and sonicated on ice for 8 min; 10s on, 10s off, 20% amplitude. The lysate was pelleted at 20k x g for 30 min, and the soluble fraction was collected. The soluble fraction was supplemented up to 15 mM imidazole and 500 mM NaCl and added to Ni-NTA agarose beads equilibrated in wash buffer (lysis buffer + 20 mM imidazole). The bead-lysate mixture was incubated for 1 h at 4 °C, rotating. Target protein was eluted in elution buffer (lysis buffer + 300 mM imidazole) and dialyzed overnight at 4 °C in dialysis buffer (20 mM HEPES, pH 8, 150 mM NaCl, 1 mM DTT) in the presence of TEV protease to cleave off the His_6_-MBP tag. Dialyzed protein was then subjected to a second Ni-NTA purification to remove the His_6_-TEV and His_6_-MBP. The flow through fraction was collected and again dialyzed overnight in dialysis buffer. The next day, the purified protein was concentrated in a 3 kDa MWCO concentrator. Protein purity was assessed by SDS-PAGE and size-exclusion chromatography. 1 mg / mL aliquots were flash frozen in liquid nitrogen and stored at -80 °C.

### Transmission electron microscopy

Quantifoil R2/2 grids coated with continuous 2 nm carbon foil were glow discharged for 45 s at 25 mA. 5 μl of protein at 0.5 mg / mL was applied to the grid for 30 s, washed 2x water, stained 30 s 2 % uranyl acetate, and dried under incandescent lamp. Grids were imaged on a FEI Morgagni microscope at 100 kV at 32,000 x magnification. For the SP110 / SP100 mixing experiment, purified aliquots of each protein were thawed and mixed at the indicated ratios for 30 min before mounting on grids.

### Cryo-EM sample preparation

Quantifoil R2/2 holey carbon copper grids (Quantifoil, Grosslöbichau, Germany) were cleaned by overnight incubation in ethyl acetate and coated with 1 nm continuous carbon foil by floatation. The grids were subjected to glow discharge (15 mA for 15 s) in a Pelco easiGlow glow discharge cleaning system (Ted Pella, Redding, CA). The climate chamber of a Vitrobot Mark IV (Thermo Fisher) was equilibrated to 4 °C at 95 % humidity. 4 μL of SP100 (crosslinked with 0.5 % glutaraldehyde and containing 0.05 % NP40) were applied to the grid, incubated for 30 s, blotted for 1 s, and subsequently vitrified by plunging into ethane-propane mixture (1:2) maintained at liquid nitrogen temperature (77 K).

### Cryo-EM data collection and image processing

10,995 movies were collected in counting mode on a Titan Krios cryo-electron microscope (Thermo Fisher) equipped with a K2 direct electron detector (Gatan, Inc.) at a nominal magnification of 165,000 x, equaling a pixel size of 0.84 Å on the specimen level. The energy filter slit width was set to 20 eV. Micrographs were exposed for 7 s and fractionated into 40 frames, with a dose rate of 8 e^-^ / px / s and an electron dose of 64 e^-^ / Å^2^. Data were collected at defocus set between -1 μm and -3 μm. Image processing was performed in cryoSPARC^66^. Movies were drift corrected, averaged, and dose weighted with Patch Motion Correction. CTF fitting was performed with Patch CTF. Filament Tracer was used for particle picking, using a 70 Å filament diameter, 0.25 x repeat distance, and 140 Å minimum length, resulting in 3,201,771 particles. The 2,884,159 particles that remained after particle picking inspection and particle extraction (360 pixels box size binned to 96 pixels) were 2D classified into 200 classes, resulting in 362,138 particles, which were subsequently extracted without binning (box size: 360 pixel). Three subsequent 2D classifications resulted in 296,140 particles that were refined to a resolution of 3.5 Å with Helix Refine, using an initial model that had been generated from a subset of 74,239 particles using Helix Refine and initial helical parameters that had been determined from the initial model using Symmetry Search. The refined helical parameters of the final reconstruction were 46.4° helical twist and 19.8 Å helical rise.

### Cryo-EM model building and refinement

An initial model of the SP100 card domain was generated with AlphaFold2 structure prediction^42^ and placed into the symmetrized, sharpened map of the SP100 filament using the rigid body fitting function “Fit in Map” of UCSF Chimera^67^. The atomic model was rebuilt in Coot^68^. The model was refined with application of non-crystallographic symmetry using Phenix real-space refinement^69^. The final structures will be available in the EMDB and PDB upon publication.

### Statistics & reproducibility

All bar graphs demonstrate the mean ± standard deviation except where noted. All biological replicates were performed at independent time points. A Pearson correlation coefficient was calculated for Figure S2h. Pearson object correlation coefficients were calculated for Figure S5f and a two-tailed Mann-Whitney U test was performed on the aggregate data.

### Software, data, and code availability

All graphs were produced in Prism (v10.1.1; GraphPad), and figures were assembled using Affinity Designer (v1.10.8; Serif). Biorender was used to prepare schematics. Sequencing data from the CRISPR screens, RNA-seq, and ChIP-seq will be deposited in an SRA BioProject prior to publication. The SP100 structure will be deposited in the EMDB and PDB prior to publication. No custom code was used in this study.

## Acknowledgements

We would like to acknowledge people at ETH Zurich facilities who aided in parts of this work: Susanne Kreutzer at Genome Engineering and Measurement Lab (GEML), Tobias Kockmann at the Functional Genomics Center Zurich (FGCZ) of University of Zurich and ETH Zurich for proteomics work, and Joachim Hehl at the Scientific Center for Optical and Electron Microscopy (ScopeM) for imaging assistance. CryoEM data were collected at ScopeM. We thank Ana Gvozdenovic for critical reading of the manuscript. EJA is supported by an EMBO Postdoctoral Fellowship (ALTF 144-2021). JEC is supported by the NOMIS Foundation and the Lotte und Adolf Hotz-Sprenger Stiftung. This project has received funding from the European Research Council (ERC) under the European Union’s Horizon 2020 research and innovation programme (855741-DDREAMM-ERC-2019-SyG) and SNSF Project Funding (310030_188858) awarded to JEC.

## Author Contributions

Project was conceived by EJA and JEC. EJA designed experiments. EJA, TK, and LS performed experiments. JR performed TEM, cryo-EM acquisition, and structural refinement with the assistance of DB. EJA and JEC wrote the manuscript with contributions from all other authors.

