## Supplementary Information for "SP110 sequestration of SP100 protects against toxic filaments during innate immune signaling"

#### Table of contents

**Figure S1**. Validation of SP110 knockdown and knockout
**Figure S2**. SP100 promotes toxicity in the absence of SP110
**Figure S3**. Few transcriptional changes between *SP110* KO and *SP110*;*SP100* KO cells
**Figure S4**. SP100 overexpression induces DNA damage and senescence
**Figure S5**. PML bodies colocalize with SP100-induced DNA damage
**Figure S6**. SP110 and SP100 localization and complementation expression
**Figure S7**. SP100 ChIP-seq extended data
**Figure S8**. Schematic representation of the cryo-EM image processing workflow
**Figure S9**. Comparison of CARD filaments
**Figure S10**. Controlling SP100 CARD filament presence
**Figure S11**. Representative flow cytometry gating strategy

**Table S1**. Cryo-EM data collection, refinement, and validation statistics
**Table S2**. Nucleic acid sequences
**Table S3**. Cell lines
**Table S4**. Antibodies
**Table S5**. Amino acid sequences

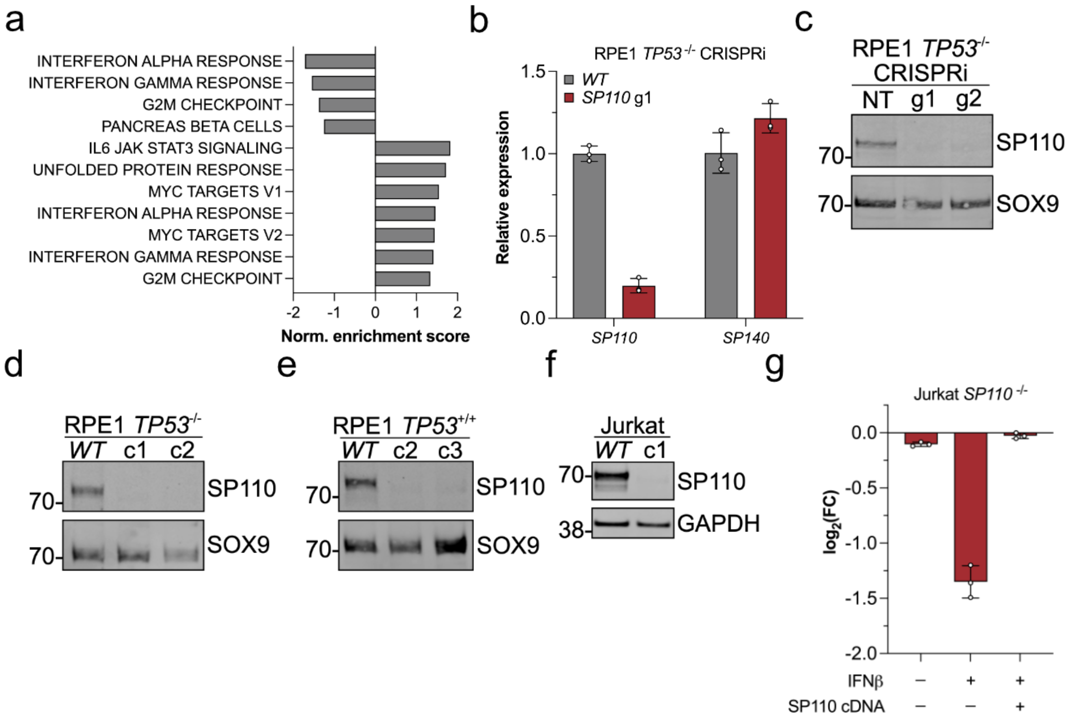

**Figure S1**. **Validation of SP110 knockdown and knockout**. **a**. Gene Set Enrichment Analysis (GSEA) from genome-wide screen from either the genes either depleted (negative normalized enrichment score) or enriched (positive normalized enrichment score) in the ppp-RNA treated condition classified according to the Molecular Signatures Database (MSigDB) hallmark gene lists with a nominative p-value $< 0.05$. **b**. Validation of *SP110* specific knockdown by CRISPRi at the RNA level. Data points are technical replicates. Data is representative of multiple independent biological replicates. **c**. Validation of SP110 specific knockdown at the protein level. NT = non-targeting guide RNA. **d**, **e**, and **f**. Validation of SP110 specific knockout of selected clones in indicated cell background. c1 = clone 1. **g**. Competition assay in Jurkat *SP110* KO cells +/- IFNβ treatment and +/- SP110 stable overexpression. Data points represent individual biological replicates. Bars in **b** and **g** denote mean +/- SD.

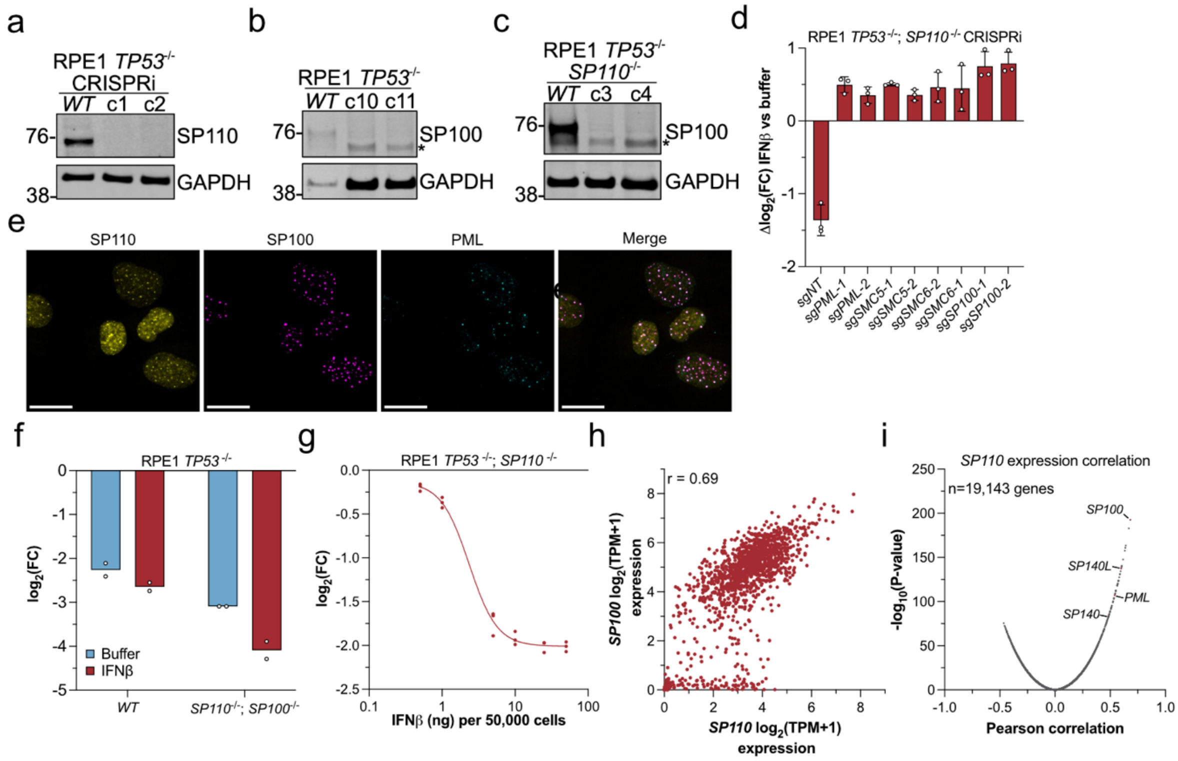

**Figure S2**. **SP100 promotes toxicity in the absence of SP110**. **a**. Confirmation of *SP110* KO clones in the RPE1 *TP53*$^{-/-}$ CRISPRi background. **b** and **c**. Western blot validation of *SP100* KO in the indicated cell types. * denotes a non-specific band. **d**. Competition assay to validate candidate genes from the CRISPRi screen in *SP110* KO cells comparing on day 7 IFNβ treatment versus buffer. Bars denote mean +/- SD. **e**. Immunofluorescence of GFP-SP110, mCherry-SP100, and PML in RPE1 *WT* cells treated with IFNβ. Scale bar = 20 μm. **f**. Competition assay in the indicated genotypes stably expressing mCherry-SP100. Bars denote mean. **g**. IFNβ titration in *SP110* KO cells. Line denotes sigmoidal 4 parameter logistic curve. **h**. SP100 versus SP110 expression across all cell lines in the Cancer Cell Line Encyclopedia (CCLE) (n = 1474 cell lines). r = Pearson correlation coefficient. **i**. Comparison of Pearson correlation coefficients between expression of SP110 and every gene (n = 19,143) in the CCLE. Data points in **d**, **f**, and **g** represent individual biological replicates.

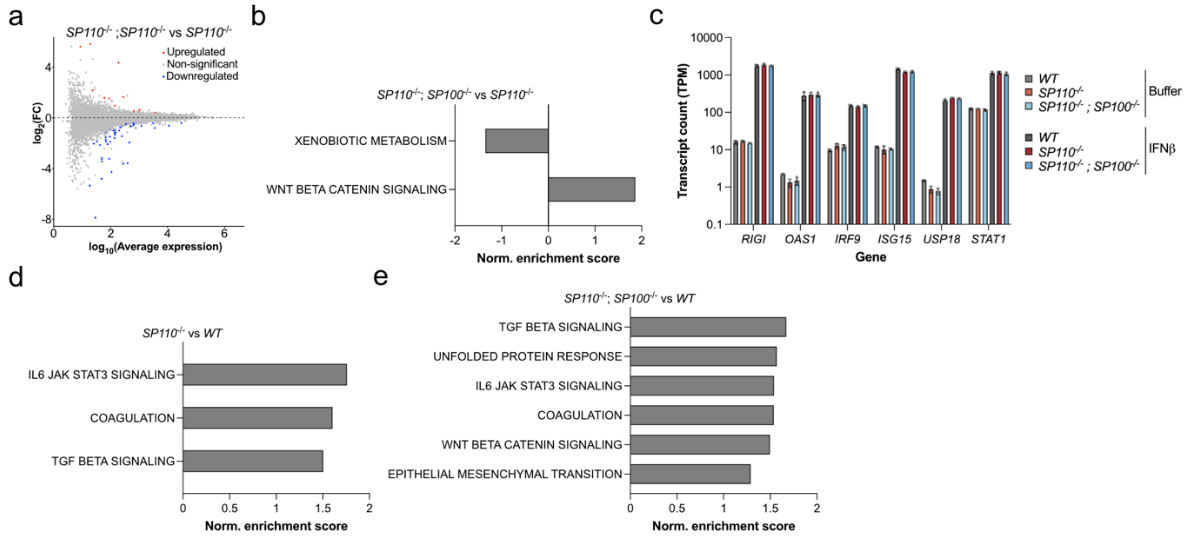

**Figure S3**. **Few transcriptional changes between *SP110* KO and *SP110*;*SP100* KO cells**. **a**. MA plot of $\log_2$ fold change (FC) in gene expression vs $\log_{10}$ average gene expression when comparing RPE1 *SP110*;*SP100* KO versus *SP110* KO cells as measured by RNA-seq under IFNβ stimulated conditions. Each data point represents a gene. Red and blue data points = adjusted p-value < 0.05. Experiment was performed with biological triplicates. **b**. Gene Set Enrichment Analysis (GSEA) classified according to the Molecular Signatures Database (MSigDB) hallmark gene lists with a nominative p-value < 0.05 for the RPE1 *SP110*;*SP100* KO versus *SP110* KO comparison under IFNβ stimulated conditions. **c**. Transcript count (in transcripts per million, TPM) of various ISGs. Each data point represents an individual biological replicate. Bars denote mean +/- SD. **d** and **e**. GSEA classified according to the MSigDB hallmark gene lists with a nominative p-value < 0.05 for the indicated genotype comparisons under IFNβ stimulated conditions.

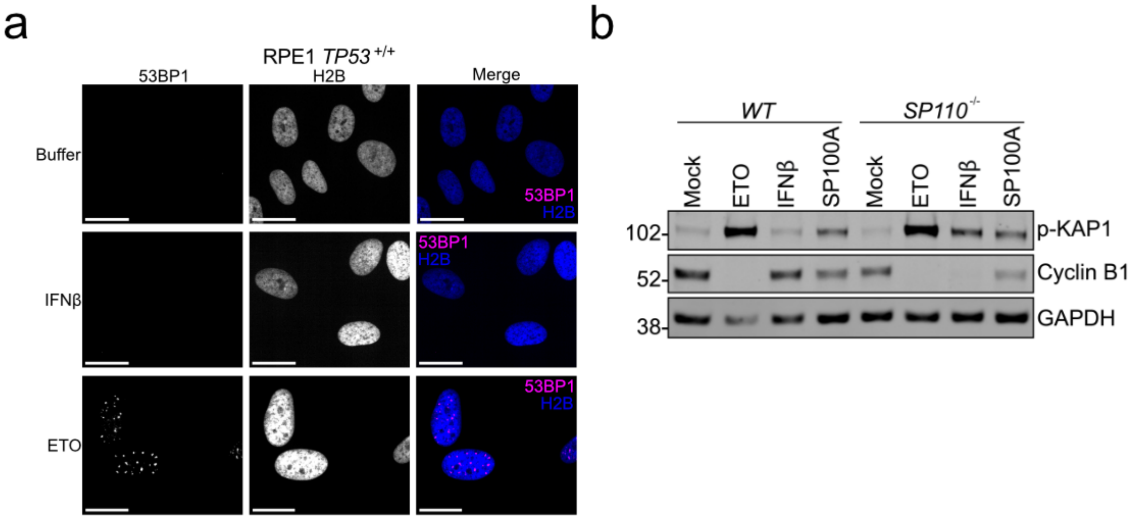

**Figure S4**. **SP100 overexpression induces DNA damage and senescence**. **a**. 53BP1 immunofluorescence in RPE1 *TP53* $^{+/+}$ *WT* cells stably expressing H2B-GFP treated with buffer, IFNβ (10 ng / mL), or etoposide (ETO; 25 μM) for 72 h. Scale bar = 20 μm. **b**. Western blot of the lysates from the indicated RPE1 *TP53* $^{+/+}$ genotypes treated with buffer (mock), ETO, IFNβ, or transduced with SP100A. Note the transduction efficiency was ~30 % and cells were not selected for positive integrations. Lysates were probed for phosphorylated KAP1 (S824) and cyclin B1 with GAPDH serving as the loading control.

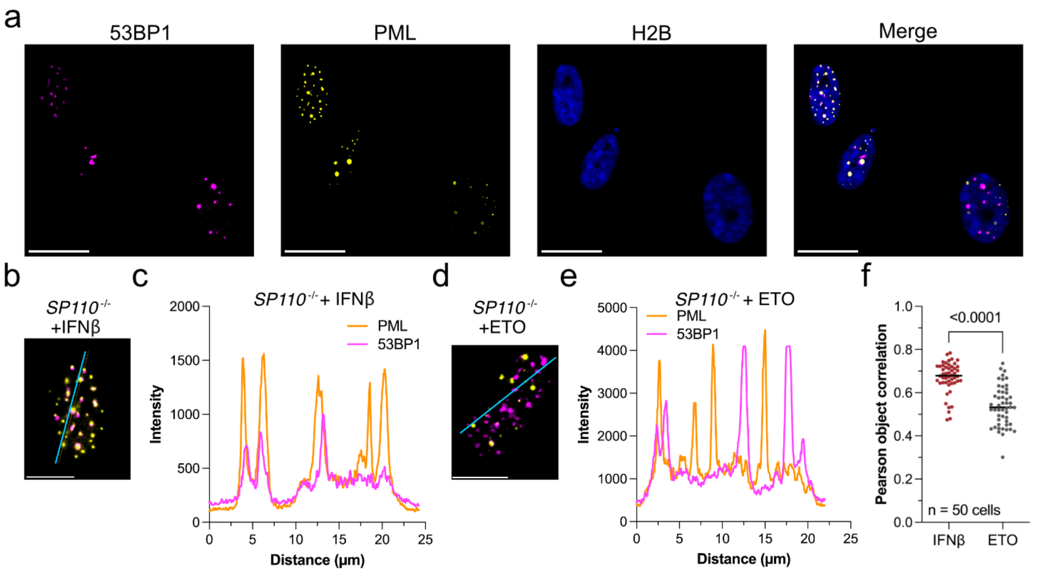

**Figure S5**. **PML bodies colocalize with SP100-induced DNA damage**. **a**. Immunofluorescence in RPE1 *SP110* KO cells stably expressing H2B-GFP stimulated with IFNβ staining for 53BP1 and PML. Scale bar = 20 µm. **b**. Individual IFNβ treated *SP110* KO cell stained with 53BP1 (magenta) and PML (yellow) with blue profile line drawn across. Scale bar = 10 µm. **c**. Quantification of intensity across the blue profile line in **b**. **d**. Individual etoposide treated *SP110* KO cell stained with 53BP1 (magenta) and PML (yellow) with blue profile line drawn across. Scale bar = 10 µm. **e**. Quantification of intensity across the blue profile line in **d**. **f**. Pearson object correlation of colocalization between PML and 53BP1 foci. Each data point represents one cell (n = 50 for each condition). A Mann-Whitney U test was used to determine statistical significance.

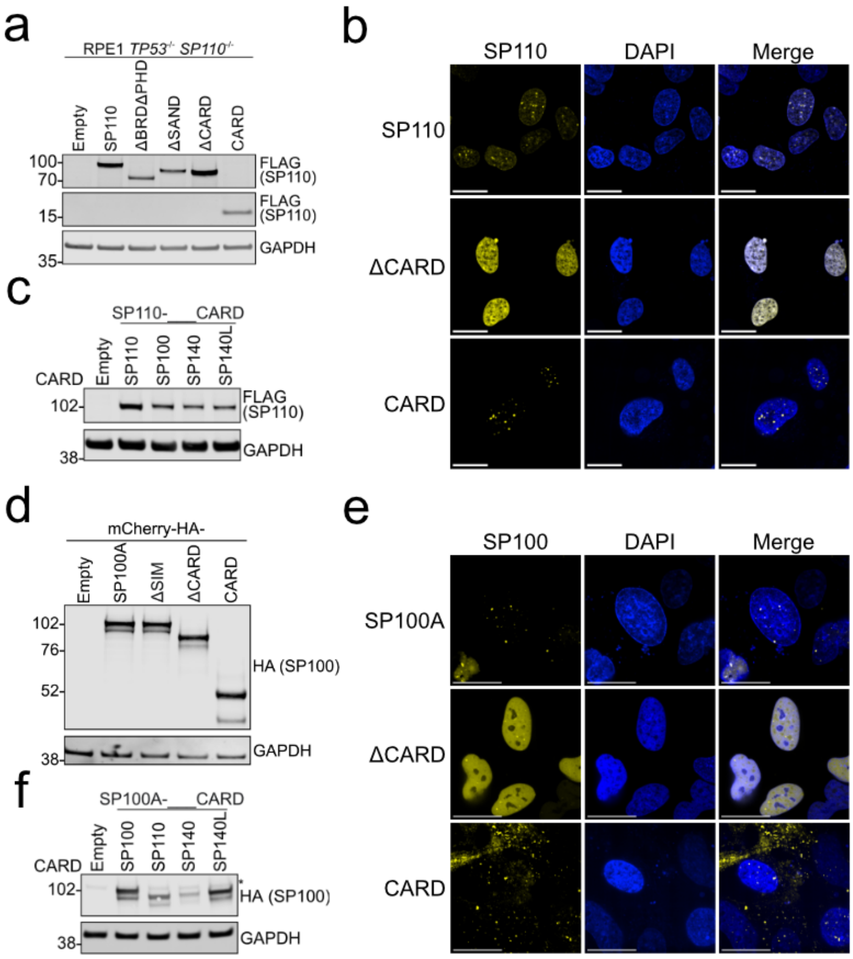

**Figure S6**. **SP110 and SP100 localization and complementation expression**. **a**. Western blot of stably expressed FLAG-SP110 variants in RPE1 *SP110* KO background. **b**. Fluorescence microscopy imaging of stably expressed SP110-GFP (yellow) constructs in RPE1 *WT* cells treated with IFNβ. Scale bar = 20 μm. **c**. Western blot of stably expressed FLAG-SP110 CARD variants RPE1 *SP110* KO background. The indicated CARD replaces the endogenous SP110 CARD in the full-length SP110. **d**. Western blot of stably expressed mCherry-HA-SP100 variants in RPE1 *SP100* KO background. **e**. Fluorescence microscopy imaging of stably expressed mCherry-SP100 (yellow) constructs in RPE1 *WT* cells treated with IFNβ. Scale bar = 20 μm. **f**. Western blot of stably expressed mCherry-HA-SP100 CARD variants RPE1 *SP100* KO background. The indicated CARD replaces the endogenous SP100 CARD in SP100A.

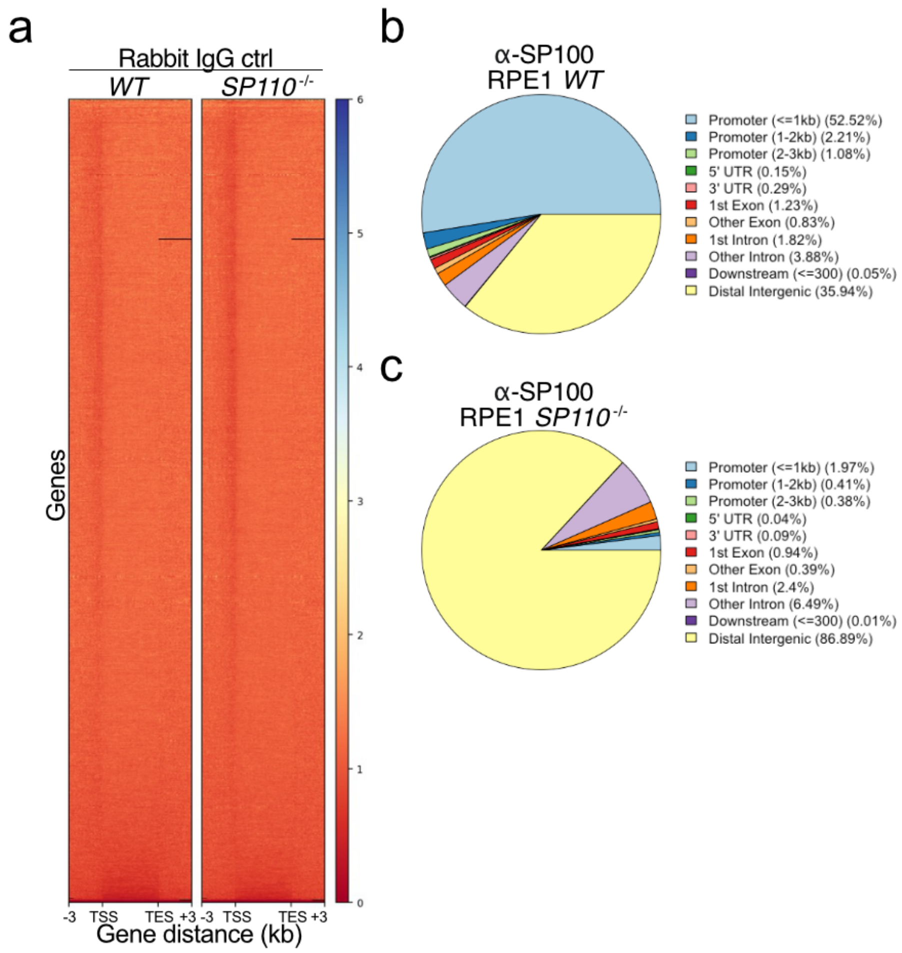

**Figure S7**. **SP100 ChIP-seq extended data**. **a**. Heat map of rabbit IgG isotype control ChIP-Seq peaks in and surrounding gene bodies of normalized length. Each line corresponds to a gene. The colour scale indicates the relative intensity of the signal. **b** and **c**. SP100 peak annotations in **b**. RPE1 *WT* background and **c**. *SP110* KO background.

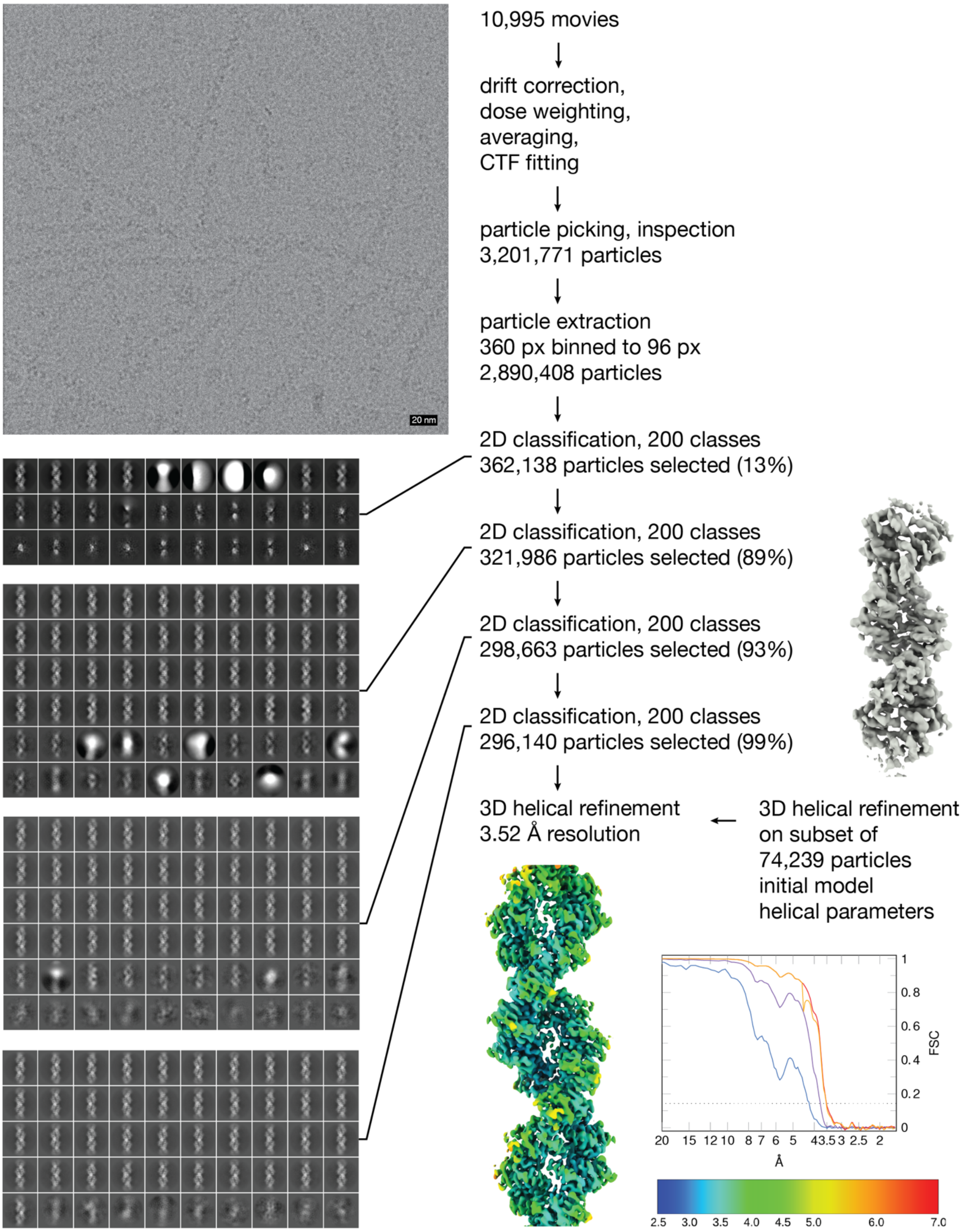

**Figure S8**. **Schematic representation of the cryo-EM image processing workflow**. Representative electron micrograph, averaged after gain correction, motion correction and dose weighting. Scale bar = 20 nm. The 2D class averages are sorted by occupancy, showing the most populated 30, 60, and 50 classes of 200 classes, respectively, with a box size of 30 nm. The FSC plot shows FSC curves without masking (blue), with loose masking (purple), tight masking (red), and corrected (yellow); correlation of 0.143 is indicated by the dotted line.

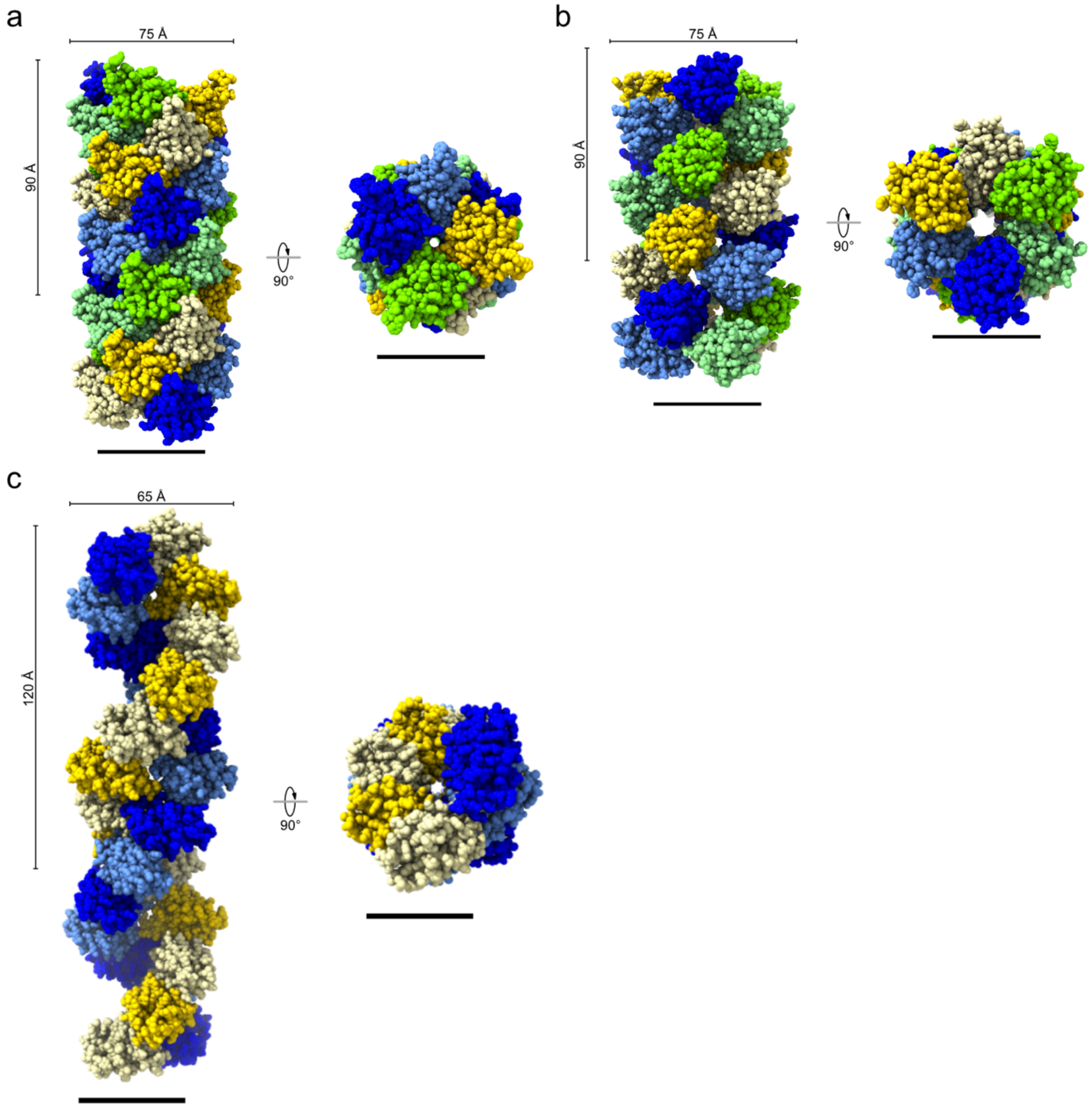

**Figure S9**. **Comparison of CARD filaments**. Cryo-EM structures of CARD filaments of **a**. NLRC4 (PDB: 6MKS), **b**. MAVS (PDB: 3J6C), and **c**. SP100 (this study). Scale bar = 50 Å. The top dimension corresponds to the filament width and the side dimension corresponds to length of one turn of the helix.

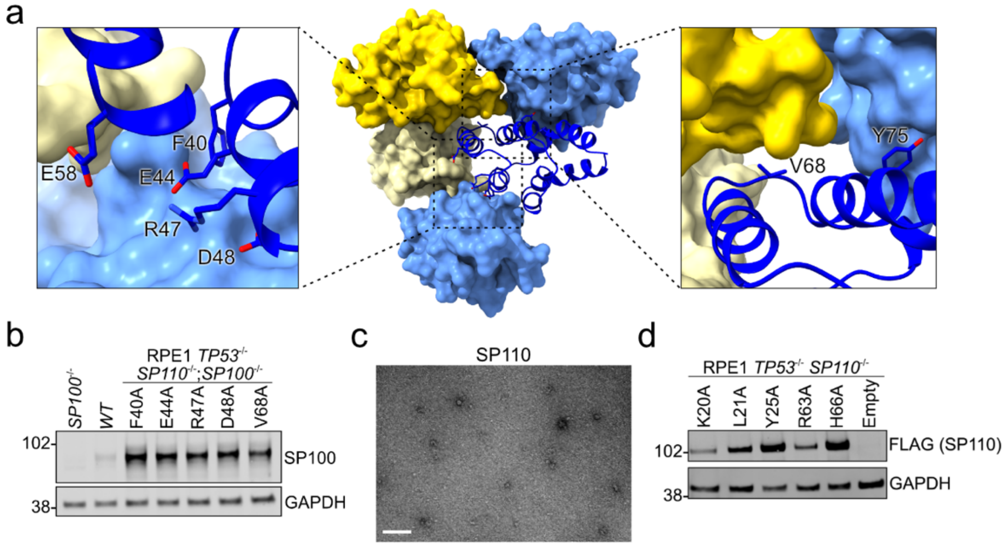

**Figure S10**. **Controlling SP100 CARD filament presence**. **a**. Detailed view of the locations of the engineered alanine point mutations in the SP100 CARD filament. **b**. Western blot of SP100 CARD point mutants stably expressed in RPE1 *SP110*;*SP100* dual KO cells. **c**. Transmission electron micrograph of recombinantly expressed SP110 CARD. **d**. Western blot of SP110 CARD point mutants stably expressed in RPE1 *SP110* KO cells.

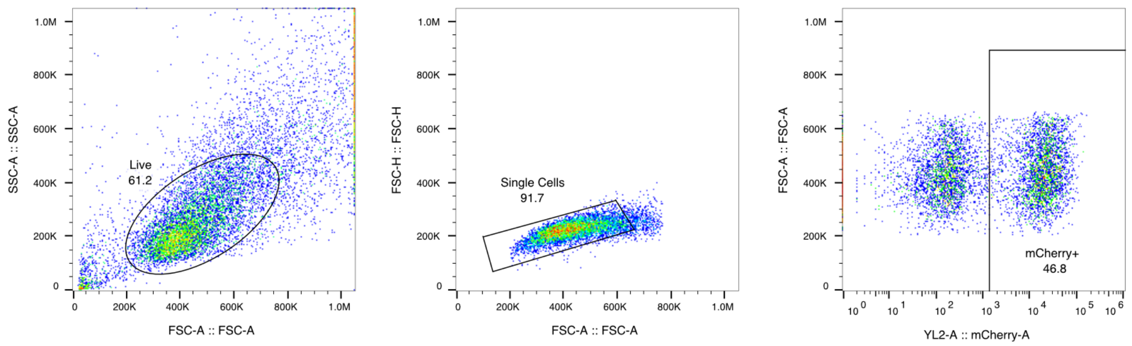

**Figure S11**. **Representative flow cytometry gating strategy**. Live cells are gated on SSC-A vs FSC-A. Single cells are gated on FSC-H vs FSC-A. Fluorescent positive cells are gated on FSC-A vs XFP (mCherry, GFP, or BFP).

**Table S1**. **Cryo-EM data collection, refinement, and validation statistics**

| | SP100 CARD filament (EMDB-19911) (PDB 9ER8) |
|---|---|
| **Data collection and processing** | |
| Magnification | 165’000x |
| Voltage (kV) | 300 |
| Electron exposure (e–/Å$^2$) | 64 |
| Defocus range (μm) | -1 to -3 |
| Pixel size (Å) | 0.84 |
| Symmetry imposed | Helical |
| Initial particle images (no.) | 2’884’159 |
| Final particle images (no.) | 296’140 |
| Map resolution (Å) FSC threshold | 3.52 (0.143) |
| Map resolution range (Å) | 1.783-46.238 |
| **Refinement** | |
| Initial model used (PDB code) | AlphaFold |
| Model resolution (Å) FSC threshold | 3.5 (0.143) |
| Model resolution range (Å) | |
| Map sharpening *B* factor (Å$^2$) | 101.8 |
| Model composition | |
| Non-hydrogen atoms | 21502 |
| Protein residues | 2574 |
| Ligands | 0 |
| *B* factors (min/max/mean; Å$^2$) | 44.95/94.16/66.53 |
| Protein | |
| Ligand | |
| R.m.s. deviations | |
| Bond lengths (Å) | 0.002 |
| Bond angles (°) | 0.388 |
| Validation | |
| MolProbity score | 0.94 |
| Clashscore | 1.81 |
| Poor rotamers (%) | 0.0 |
| Ramachandran plot | |
| Favored (%) | 98.97 |
| Allowed (%) | 1.03 |
| Disallowed (%) | 0 |

### Table S2. Nucleic acid sequences

#### ppp-RNA sequence

| Name | Sequence (5'-to-3') |
|---|---|
| ppp-RNA | GCCAAGCTGAAGGTGACCAAGTTTTAGAGCTAGAAATAGCAAGTTAAAATAA GGCTAGTCCGTTATCAACTTGAAAAAGTGGCACCGAGTCGGTGCTTTTTTT |

#### sgRNA sequences

| Name | Spacer sequence (5'-to-3') |
|---|---|
| *sgSP110-1* KD | GGGGATCACTCCTCAAGATT |
| *sgSP110-2* KD | GCAAGATTGGGAGAGTTACA |
| *sgSP100-1* KD | GGGCCGACTGAGGGGCTCAG |
| *sgSP100-2* KD | GGCAGGCTGGGCCGACTGAG |
| *sgPML-1* KD | GGTGCGTGAAGAGAAGCTGA |
| *sgPML-2* KD | GAGTAGAAAAGAGGACACGG |
| *sgZNFX1-1* KD | GAAGCCCCAGCACTCGCCGG |
| *sgZNFX1-2* KD | GGCGAGGCAAGGAAGAATCA |
| *sgCNOT2-1* KD | GAGCGGCGGTAAGGGCGGTA |
| *sgCNOT2-2* KD | GCGGGACAAGAAAATTCATG |
| *sgUSP18-1 KD* | GCGAGCGCGGGGCCAGACCA |
| *sgUSP18-2 KD* | GACGTGGAACTCAGCAGCGG |
| *sgSMC5-1* KD | GGCCCGGACGACTGCAACAG |
| *sgSMC5-2* KD | GCAGCACACCTCCACACCCG |
| *sgSMC6-1* KD | GCAGGAGAAGGTAGATTCCC |
| *sgSMC6-2* KD | GGCAGGCGCGGTTAGTACCG |
| *sgSP110* KO | CCAGATTGGGATATTCACGC |
| *sgSP100* KO | GCAGCCTGTCATCTACACCC |
| *sgNT* | GCGGTGGGGCCGCTCACTAG |

#### PCR primers

| Name | Forward primer (5'-to-3') | Reverse primer (5'-to-3') |
|---|---|---|
| *SP110* NGS | CTTTCCCTACACGACGCTCTTCCGATCT TCTCACCCAACTGGAGAGGA | GGAGTTCAGACGTGTGCTCTTCCGATCT CCACAGGGCTAGAAGTAAACC |
| *SP100* NGS | CTTTCCCTACACGACGCTCTTCCGATCT TATGTAAACTCTTACAGCCTCC | GGAGTTCAGACGTGTGCTCTTCCGATCT GGAGGCCCTCGAGGAATGGAA |
| *ACTB* qPCR | GGGTCAGAAGGATTCCTATG | GGTCTCAAACATGATCTGGG |
| *SP110* qPCR | GGGACAGCCTCATCTAGACA | AAGTGCCATCAATGATCTCCTC |
| *SP100* qPCR | GGAGAAGAGCTTCAGGAAACCTG | GGCTTCTTGGCACACCTTTTGG |
| *SP140* qPCR | GCATGGCTGGAGCAGAATGAGA | CGTTCGCCTTCTTTTCACTGGAG |
| *CDKN1A (p21)* qPCR | AGGTGGACCTGGAGACTCTCAG | TCCTCTTGGAGAAGATCAGCCG |

**Table S3. Cell lines**

| **Cell line** | **Source** |
|---|---|
| HEK-293T | ATCC |
| RPE1 *TP53* $^{-/-}$ | Dr. Steve Jackson |
| RPE1 *TP53* $^{+/+}$ | ATCC |
| RPE1-dCas9-BFP-KRAB(KOX1) *TP53* $^{-/-}$ | Fielden, Siegner, et al. 2023 |
| RPE1-dCas9-BFP-KRAB(KOX1) *SP110* $^{-/-}$; *TP53* $^{-/-}$ | This study |
| RPE1 *TP53* $^{-/-}$; *SP110* $^{-/-}$ | This study |
| RPE1 *TP53* $^{+/+}$; *SP110* $^{-/-}$ | This study |
| RPE1 *TP53* $^{-/-}$; *SP100* $^{-/-}$ | This study |
| RPE1 *TP53* $^{-/-}$; *SP110* $^{-/-}$; *SP100* $^{-/-}$ | This study |
| RPE1 *TP53* $^{+/+}$; *SP110* $^{-/-}$; *SP100* $^{-/-}$ | This study |
| RPE1 *TP53* $^{+/+}$ H2B-GFP | This study |
| RPE1 *TP53* $^{+/+}$; *SP110* $^{-/-}$ H2B-GFP | This study |
| Jurkat | ATCC |
| Jurkat *SP110* $^{-/-}$ | This study |
| Jurkat *SP100* $^{-/-}$ | This study |
| Jurkat *SP110* $^{-/-}$; *SP100* $^{-/-}$ | This study |

### Table S4. Antibodies

| Primary Antibody | Application | Species | Supplier | Product # | Clone # | Lot # | Working dilution |
|---|---|---|---|---|---|---|---|
| SP110 | WB | Ms | Santa Cruz Biotech | sc-376741 | Monoclonal B-10 | K0116 | 1:1000 |
| SP100 | WB, IF, ChIP | Rb | Novus Biologicals | NBP1-86060 | Polyclonal | A106767 | 1:1000 (WB), 1:200 (IF) |
| FLAG | WB, IP | Ms | Sigma | F3165 | Monoclonal M2 | SLCQ9255 | 1:1000 (WB) |
| HA | WB | Rb | Cell Signalling Technologies | 3724 | Monoclonal C29F4 | 11 | 1:1000 (WB) |
| P-KAP1 (S824) | WB | Rb | Abcam | ab70369 | Polyclonal | GR3316128-13 | 1:1000 |
| Cyclin B1 | WB | Rb | R&D Systems | MAB60001 | Monoclonal 2061D | CKJC022109A | 1:1000 |
| GAPDH | WB | Rb | Cell Signalling Technologies | 2118 | Monoclonal 14C10 | 16 | 1:2000 |
| GAPDH | WB | Ms | Cell Signalling Technologies | 97166 | Monoclonal D4C6R | 7 | 1:2000 |
| SOX9 | WB | Rb | Abcam | ab185966 | Monoclonal EPR14335-78 | GR3183675-1 | 1:1000 |
| 53BP1 | IF | Rb | Novus Biologicals | NB100-304 | Polyclonal | D148226 | 1:1000 |
| PML | IF | Ms | Santa Cruz | sc-966 | Monoclonal PG-M3 | G1122 | 1:200 |
| Rabbit IgG isotype control | ChIP | Rb | Novus Biologicals | NBP2-24891 | Polyclonal | n/a | |

| Secondary Antibody | Application | Species | Supplier | Product # | Clone # | Lot # | Working dilution |
|---|---|---|---|---|---|---|---|
| IRDye 800CW anti-Rabbit IgG | WB | Gt | LI-COR Biosciences | 926-32213 | Polyclonal | D30207-15 | 1:10000 |
| IRDye 800CW anti-Mouse IgG | WB | Gt | LI-COR Biosciences | 926-32212 | Polyclonal | D30124-05 | 1:10000 |
| IRDye 680RD anti-Rabbit IgG | WB | Gt | LI-COR Biosciences | 926-68073 | Polyclonal | D30328-05 | 1:10000 |
| IRDye 680RD anti-Mouse IgG | WB | Gt | LI-COR Biosciences | 926-68072 | Polyclonal | D20503-05 | 1:10000 |
| Anti-rabbit Alexa Fluor Plus 405 | IF | Gt | Invitrogen | A48254 | Polyclonal | YD369487 | 1:10000 |
| Anti-mouse Alexa Fluor 568 | IF | Gt | Invitrogen | A11004 | Polyclonal | 2447869 | 1:10000 |

#### Table S5. Amino acid sequences

| Construct | Amino acid sequence (N-to-C) |
|---|---|
| His$_6$-MBP-TEV-SP100 CARD | MKSSHHHHHHGSSMKIEEGKLVIWINGDKGYNGLAEVGKKFEKDTGIKVTVEHPDK<br>LEEKFPQVAATGDGPDIIFWAHDRFGGYAQSGLLAEITPDKAFQDKLYPFTWDAVR<br>YNGKLIAYPIAVEALSLIYNKDLLPNPPKTWEEIPALDKELKAKGKSALMFNLQEPYFT<br>WPLIAADGGYAFKYENGKYDIKDVGVDNAGAKAGLTFLVDLIKNKHMNADTDYSIAE<br>AAFNKGETAMTINGPWAWSNIDTSKVNYGVTVLPTFKGQPSKPFVGVLSAGINAAS<br>PNKELAKEFLENYLLTDEGLEAVNKDKPLGAVALKSYEEELAKDPRIAATMENAQKG<br>EIMPNIPQMSAFWYAVRTAVINAASGRQTVDEALKDAQTNSSSNNNNNNNNNNLGI<br>EENLYFQSNATDLQRMFTEDQGVDDRLLYDIVFKHFKRNKVEISNAIKKTFPFLEGL<br>RDRDLITNKMFEDSQDSCRNLVPVQRVVYNVLSELEKTFNLPVLEALFSDVNMQEY<br>PDLIHIYKGFENVIH |
| His$_6$-MBP-TEV-SP110 CARD | MKSSHHHHHHGSSMKIEEGKLVIWINGDKGYNGLAEVGKKFEKDTGIKVTVEHPDK<br>LEEKFPQVAATGDGPDIIFWAHDRFGGYAQSGLLAEITPDKAFQDKLYPFTWDAVR<br>YNGKLIAYPIAVEALSLIYNKDLLPNPPKTWEEIPALDKELKAKGKSALMFNLQEPYFT<br>WPLIAADGGYAFKYENGKYDIKDVGVDNAGAKAGLTFLVDLIKNKHMNADTDYSIAE<br>AAFNKGETAMTINGPWAWSNIDTSKVNYGVTVLPTFKGQPSKPFVGVLSAGINAAS<br>PNKELAKEFLENYLLTDEGLEAVNKDKPLGAVALKSYEEELAKDPRIAATMENAQKG<br>EIMPNIPQMSAFWYAVRTAVINAASGRQTVDEALKDAQTNSSSNNNNNNNNNNLGI<br>EENLYFQSNATMEEALFQHFMHQKLGIAYAIHKPFPFFEGLLDNSIITKRMYMESLEA<br>CRNLIPVSRVVHNILTQLERTFNLSLLVTLFSQINLREYPNLVTIYRSFKRV |
